# MIF-induced CD74+ microglia/macrophages are clinically relevant disease-associated subpopulations in brain metastasis and other CNS disorders

**DOI:** 10.1101/2025.09.06.673623

**Authors:** Laura Álvaro-Espinosa, Angel Marquez-Galera, Neibla Priego, Virginia García-Calvo, María Perea-García, Carolina Hernández-Oliver, Diana Retana, Oliva Sánchez, Ana de Pablos-Aragoneses, Pedro García-Gómez, Osvaldo Graña-Castro, Óscar Lapuente-Santana, Fátima Al-Shahrour, Ana Cayuela, Isabel Peset, Diego Megías, Mihaela Ola, Damir Varešlija, Leonie Young, Yolanda Martí-Mateos, José Antonio Enríquez, Elena Hernández-Encinas, Carmen Blanco-Aparicio, Marisol Soengas, Jürgen Bernhagen, Alejandro Antón-Fernández, Jesús Ávila, Miguel A Marchena, Maximiliano Torres, Fernando de Castro, Mar Márquez-Ropero, Amanda Sierra, Jose P Lopez-Atalaya, RENACER, Manuel Valiente

**Author notes:** A list of authors appears at the end of the paper. Current address: Diego Megías, Advanced Optical Microscopy Unit, UCCTs, Instituto de Salud Carlos III (ISCIII), Madrid, Spain., Elena Hernández-Encinas, Dosimetría de Radiaciones Ionizantes, CIEMAT, Madrid, Spain., Alejandro Antón-Fernández, Department of Neuroscience and Biomedical Sciences, Carlos III University (UC3M), Madrid, Spain, Miguel A Marchena, NeuroLab – Faculty of Health Sciences – HM Hospitals, Camilo Jose Cela University and HM Hospitals Health Research Institute, Madrid, Spain., Maximiliano Torres, Laboratory of Neurodegeneration and Neuroprotection Mechanisms, Clemente Estable Institute of Biological Research, Montevideo, Uruguay.

## Abstract

The upregulation of CD74, a chaperone involved in MHC-II antigen processing ^1,2^, has been broadly reported in virtually all brain disorders analyzed by single-cell RNA sequencing ^3–6^. However, its expression is usually interpreted as indicative of antigen presentation. In parallel, CD74 expression has also been described in cancer cells across multiple tumor types, but interestingly in glioma its expression has been mainly identified in the microenvironment. However, the functional contribution of CD74 to disease progression in the brain, and specifically in secondary brain tumors, has not been directly addressed. Here we described that, in contrast to what it has been assumed, the presence of CD74^+^ microglia/macrophages, which is induced by increased levels of interferon gamma in the brain affected by metastases, does not relate to its canonical pathway. Instead, CD74’s alternative function as cytokine receptor is pivotal. Rewired by increasing levels of its ligand MIF, produced by proliferating cancer cells, the CD74 receptor, upon binding to this ligand, translocates to the nucleus activating a NF-κB-dependent program promoting metastasis progression. A brain metastasis-associated CD74 signature involves a more aggressive progression of the local disease in patients, while it has no clinical correlation with the matched primary tumor. Furthermore, we identified the CD74^+^ myeloid population in additional brain disorders including Alzheimer’s disease and multiple sclerosis, which shared a pan-disease non-canonical signature with clinical relevance. The brain-penetrant drug ibudilast, which prevents the binding of MIF to CD74, decreases brain metastases in experimental models *in vivo* and in patient-derived organotypic cultures *ex vivo* in a primary tumor-agnostic manner. Our findings suggest that MIF/CD74-induced reprogramming of myeloid cells in brain disorders is a novel vulnerability that could be exploited therapeutically against brain metastases, and possibly other brain disorders, guided by a non-invasive molecular strategy.

## Introduction

Secondary brain tumors are a frequent progression of cancer that involves a decreased survival and quality of life despite multimodal treatment approaches, which, however, remain palliative in most patients ^7,8^. Whether brain metastases should be treated like their primary tumors is still debated, as brain relapses are increasingly common even in patients who respond well to systemic therapies that work in both sites for certain cancer types ^9–15^. Additionally, new strategies are emerging to target brain-specific vulnerabilities mainly focused on the microenvironment ^16^. One such vulnerability is the unique vasculature of the brain (blood brain barrier, BBB), which requires specific molecular strategies that metastatic cells must activate to extravasate. For instance, targeting cathepsin S, which promotes breakage of BBB-junctions, prevents brain metastases ^17^. After completing extravasation, metastatic cells must prevent brain-specific responses reacting to the presence of micrometastases ^18^. With time, the increasing metastatic load influences the brain, inducing altered molecular patterns in subpopulations of reactive astrocytes that promote brain metastasis progression ^19^. This occurs through their role in blocking the CD8^+^ T cells infiltrating from the periphery ^20^ as well as by preventing the expression of MHC-I in cancer cells ^21^. Drugs targeting some of these mechanisms are in clinical trials (NCT05689619). One of the potential benefits of exploiting brain-specific vulnerabilities is the possibility that they could be expanded to other disorders affecting this organ. Given the high demand of effective therapeutic strategies for brain disorders in general, the identification of a pan-disease target based on the shared components of the microenvironment is pressing but remains unexplored.

Here we show that the expected anti-tumor nature of microglia/macrophages positive for CD74 (also known as the invariant chain), due to its canonical involvement in antigen processing through the MHC-II complex ^22^, is rewired into a non-canonical, pro-metastatic program in the brain metastatic niche. CD74, beyond its classical antigen-presenting role, is also a receptor for the cytokine macrophage migration inhibitory factor (MIF), a pro-inflammatory molecule involved in various cell processes, including cell survival, proliferation and migration ^23^. Upon MIF binding, CD74 can undergo endocytosis, a process that involves regulated intramembrane proteolysis, releasing its intracellular domain (ICD), which translocates into the nucleus to activate NF-κB–dependent transcription ^24–27^. This signaling cascade, previously described in B cells, is shown here to reprogram microglia/macrophages into a metastasis-supporting state in brain metastases.

It has been reported that MIF levels are highly elevated in tumors ^28^, including glioblastoma, where it can act as a dual driver of tumor progression: acting within tumor cells to promote growth and survival, while simultaneously remodeling the microenvironment to favor immune evasion, myeloid recruitment, and angiogenesis^29^. The MIF–CD74 axis has been implicated in cancer progression across multiple tumor types, including melanoma ^30^, NSCLC ^31^, pancreatic cancer ^32^ or cervical squamous cell carcinoma ^33^. However, most studies focused on CD74 expression by cancer cells themselves ^30,31,34,35^, not on immune cells within the tumor microenvironment. In several tumor types, including breast and melanoma, CD74 expression is enriched in stromal compartments such as macrophages, dendritic cells, and endothelial cells, often correlating with immune suppression and tumor progression ^36,37^. In glioma, CD74 is expressed primarily by glioma-associated microglia/macrophages (GAMs), with conflicting roles, some studies reporting anti-tumoral effects ^38^, and others linking CD74 to immune evasion and M2 polarization ^39,40^. However, secondary brain tumors represent a distinct entity with a more complex and heterogeneous immune microenvironment. Our previous work identified a pro-tumoral role for MIF–CD74 signaling in microglia/macrophages associated with brain metastases ^19^. More recently, MIF–CD74 inhibition has been shown to promote M1 polarization and enhance radiotherapy response in brain metastases from non-small cell lung cancer ^41^, although the study did not address the role of CD74^+^ population in disease progression.

Building on this previous knowledge, our current study characterizes a unique CD74⁺ subpopulation of microglia/macrophages in brain metastases with distinct transcriptional features and pro-metastatic behavior that correlate with clinical outcome in patients. We show how this population emerges during metastatic colonization of the brain after being functionally rewired by tumor-derived signals and sustained by mitochondrial plasticity, a known mechanism of macrophage adaptation^42^. Remarkably, the CD74^+^ derived transcriptomic signature we established here could be expanded to other neurodegenerative and neuroinflammatory brain disorders both in preclinical models and patient derived datasets.

The MIF-CD74 axis may be targeted by the small molecule drug ibudilast, which binds to and prevents the activity of MIF through CD74 ^43^. Ibudilast has been previously shown to penetrate the BBB and demonstrated a safe profile ^44^. Beyond its activity targeting MIF, ibudilast has also been shown to target phosphodiesterase’s involvement in inflammation ^45^. Consequently, the use of ibudilast requires strategies to assess its on target effect. Currently, ibudilast is under investigation in combination with temozolomide for newly diagnosed and recurrent glioblastoma patients (NCT03782415) ^46^, and for the treatment of multiple sclerosis (NCT01982942) ^47^, highlighting its potential role in targeting CD74-mediated pathways in the brain. In our study, we show that ibudilast impairs the MIF-CD74-dependent transcriptional program in CD74⁺ microglia/macrophages, reversing their pro-metastatic phenotype and restoring immune surveillance. This effect was validated both in preclinical mouse models and in patient-derived organotypic cultures from a range of primary tumor types, including lung, breast, and melanoma.

Together, our findings reveal a brain-specific immune vulnerability centered on the MIF–CD74 axis in microglia/macrophages, offering a tractable therapeutic strategy for targeting the tumor microenvironment in secondary brain tumors and other brain disorders. The accompanied strategies providing the possibility to use a non-invasive liquid biopsy biomarker as well as to identify patients responding to ibudilast based on a reduced transcriptomic signature support the feasibility for clinical translation.

## Results

### MIF reprograms brain metastasis-associated macrophages/microglia

We found that brain metastases, both experimental and patient samples, show high levels of MIF expressed by cancer cells (Fig. 1a, Supplementary Fig. 1a-e, Table 1). Moreover, the MIF receptor was found to colocalize with microglia/macrophage markers in experimental brain metastases and patient samples (Fig. 1b-d). Notably, CD74⁺ cells are absent from the healthy brain parenchyma, making it the only organ without basal CD74 expression under homeostatic conditions (Supplementary Fig. 1f). Thus, the emergence of CD74^+^ microglia/macrophages correlates with the local progression of brain metastases, as this cell subpopulation is still absent at early stages during metastatic colonization of the brain, where only micrometastases could be found, but highly abundant when metastases are established (Supplementary Fig. 1g). In contrast, MIF is already highly expressed by cancer cells during the early stages of brain colonization (Supplementary Fig 1h). The main inducer of *CD74* expression is interferon-gamma (IFN-γ), which activates the *Ciita* transcription factor ^48–50^. We confirmed increased IFN-γ levels during metastatic colonization of the brain (Supplementary Fig. 1i, j) and its ability to induce CD74 in resident microglia and bone marrow-derived macrophages when evaluated in an *ex vivo* system (Supplementary Fig. 1k-o). Overall, we found that MIF secretion from cancer cells and the presence of its receptor CD74 in microglia/macrophages is a common finding in mouse and human brain metastases irrespectively of the primary source.

**Figure 1.**
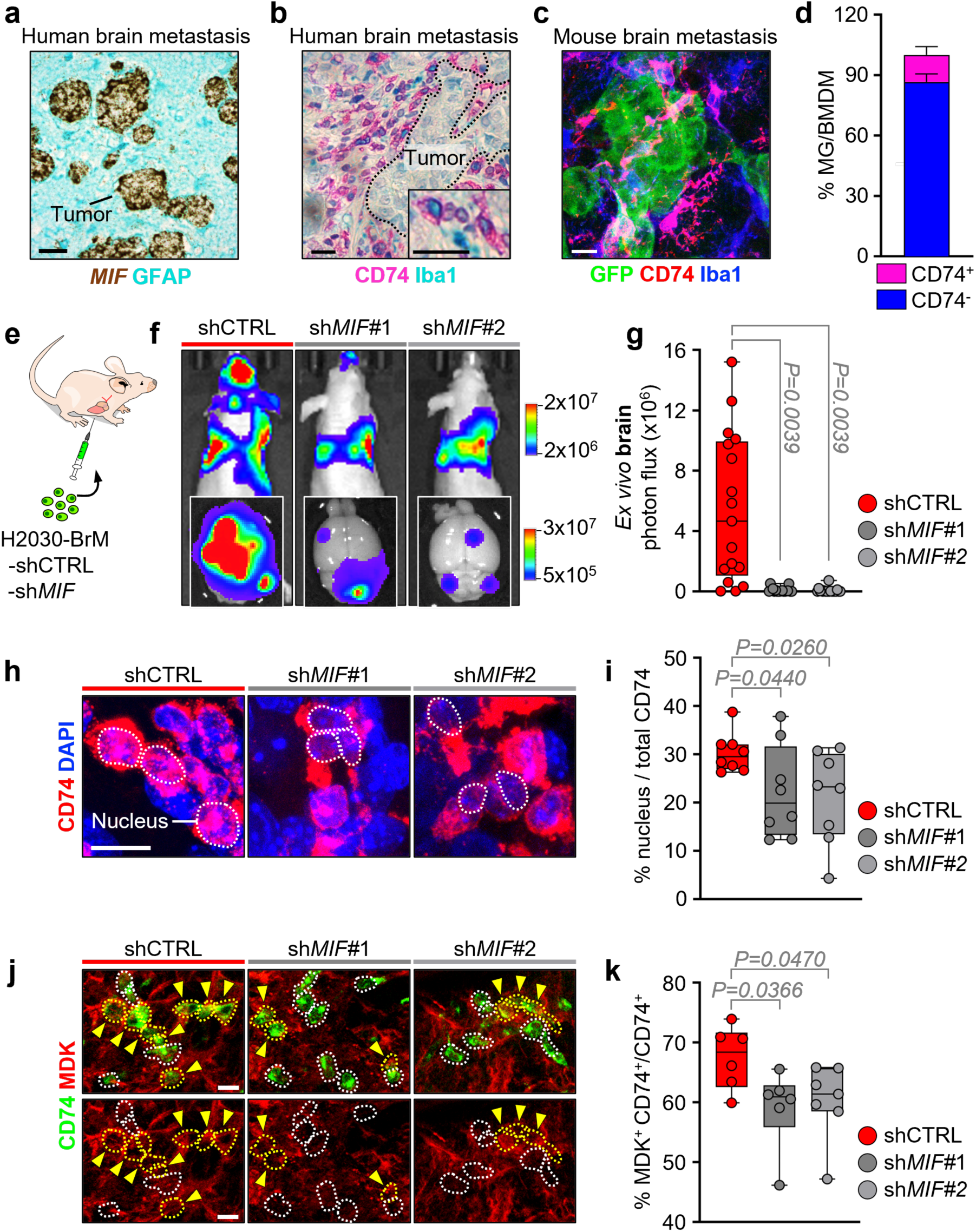
MIF as a reprogramming factor for brain metastasis macrophages/microglia. **a**. Representative image of human brain metastasis with RNA *in situ* hybridization using RNAScope against *MIF* and GFAP immunohistochemistry. Scale bar: 20 µm. **b.** Immunohistochemistry against CD74 and Iba1 in human brain metastasis. The black dotted lines delineate the tumoral area. Scale bar: 10 μm. **c.** Immunofluorescence against CD74, Iba1 and GFP (H2030-BrM) in mouse established brain metastasis. Scale bar: 10 μm. **d.** Quantification by immunofluorescence of CD74+ Iba1+ cells, which are microglia or bone marrow-derived macrophages (MG, BMDM), in H2030-BrM experimental brain metastasis model. n=6 brains, with 10 FOV per mouse, from 2 independent experiments. Bar graph represents the mean+SD. **e.** Schema of the experimental design. H2030-BrM cells harboring a knockdown (KD) for *MIF* using two different shRNAs were intracardially injected in immunocompromised wild-type mice. **f.** Bioluminescence imaging (BLI) *in vivo* of mice injected with either H2030-BrM WT cells or KD for *MIF* at endpoint (32-35 days after intracardiac injection). *Ex vivo* BLI of brains (down) are also shown. **g.** Quantification of *ex vivo* BLI of brains from f. Values are shown in a box-and-whisker plot where each dot represents a different animal and the line corresponds to the median. n=17 (shCTRL), n=18 (*shMIF#1*), n=19 (*shMIF#2*) mice from 2 independent experiments. *P* value was calculated using an unpaired two-tailed t-test of each experimental condition against control. **h.** Immunofluorescence images of CD74 and DAPI in H2030-BrM tumors. White dotted lines outline de DAPI^+^ nuclei. Scale bar: 10 µm. **i.** Quantification of CD74 found in the nucleus (DAPI) vs the membrane/cytoplasm on images shown in h. Values are shown in a box-and-whisker plot where each dot represents the mean percentage of CD74-ICD found in the nuclei per animal, and the line corresponds to the median. Quantification was done in Iba1^+^ cells. 3-12 fields of view (FOV) were captured per brain, in n=8 animals from each experimental condition, from 2 independent experiments. *P* value was calculated using an unpaired two-tailed t-test of each experimental condition against control. **j.** Immunofluorescence against CD74 and MDK H2030-BrM in tumors. White dotted lines outline the CD74^+^ MDK^−^ cells, while yellow dotted lines indicated with an arrow delineate the double-positive CD74^+^ MDK^+^ cells. Images with the red-channel only (MDK) are shown for clarity. Evaluation of the expression was done in Iba1^+^ cells. Scale bar: 10 µm. **k.** Quantification of images shown in j, represented as the mean percentage of CD74^+^ microglia/macrophages (Iba1^+^) positive for MDK normalized to the total CD74^+^ microglia/macrophages (Iba1^+^). Values are shown in a box and whisker plot where each dot represents the mean percentage of CD74^+^ MDK^+^ cells per animal. 3-12 FOV were captured per brain, in n=6 brains from each experimental condition from 2 independent experiments. *P* value was calculated using an unpaired two-tailed t-test of each experimental condition against control.

Independently of its role as an MHC-II chaperone, CD74 can act as a cytokine receptor. Upon binding to MIF, CD74 can undergo endocytosis. The processing involves cleavage, which generates an intracellular domain (ICD). This CD74-ICD can translocate to the nucleus, where it can modulate gene expression ^26,27^. To test whether MIF influences CD74^+^ microglia/macrophages, we first treated *ex vivo* control metastasis-naïve brain slices with IFN-γ alone or IFN-γ and MIF (Supplementary Fig. 2a). According to existing bibliography ^24–27,48–50^, only when both cytokines were combined nuclear translocation of CD74-ICD could be observed (Supplementary Fig. 2b, c).

To functionally evaluate the contribution of MIF in brain metastasis progression we induced loss of function of MIF in two metastatic cell lines from lung adenocarcinoma ^51^ and melanoma ^19^ including brain tropism (Fig. 1e, Supplementary Fig. 2d-f). While cancer cells without MIF grew similarly *in vitro* (Supplementary Fig. 2g), *in vivo* brain metastasis assays revealed significantly reduced tumor viability as evaluated by bioluminescence imaging (BLI) (Fig. 1f, g and Supplementary Fig. 2h-k). When systemic inoculation was used (intracardiac, iC), a significant reduction was also observed in lung metastases (Supplementary Fig. 2l-o), but the effect was notably less pronounced than in the brain (Fig. 1f-g and Supplementary Fig. 2j-k), reinforcing the brain-specific relevance of this phenotype. This reduction in brain metastasis was accompanied by decreased nuclear accumulation of CD74-ICD in metastasis-associated microglia/macrophages (Fig. 1h, i). One of the downstream effects of MIF binding to CD74 is the activation of the NF-κB pathway, leading to the transcription of genes involved in cell survival and inflammation ^24,25^. Among these genes is *midkine* (*MDK*) ^52^, an heparing-binding growth factor that plays a crucial role in cancer progression. MDK has been shown to contribute to metastatic progression by promoting cell survival, proliferation and migration of cancer cells ^53^. We also confirmed reduced expression of the pro-tumoral NF-κB-dependent gene MDK in the CD74^+^ microglia/macrophages upon MIF knockdown in the cancer cells (Fig. 1j, k).

Our data suggests that the broad MIF expression found in experimental and human brain metastases might be required to rewire the brain microenvironment by remodeling an interferon-mediated response in microglia/macrophages into a pro-metastatic CD74^+^ subpopulation.

### A brain-specific poor prognosis signature derived from CD74^+^ macrophages/microglia

Our functional data suggest the potential contribution of a disease-associated population of microglia/macrophages characterized by the expression of CD74. To evaluate its contribution to the human disease we scored whether a transcriptomic signature derived from experimental brain metastases-associated CD74^+^ microglia/macrophages could correlate with clinical parameters. We first interrogated bulk RNA-seq of Cx3cr1^+^ Cd11b^+^ CD74^+^ microglia/macrophages in comparison with the CD74-negative counterparts in two brain metastasis experimental models (Fig. 2a, Supplementary Fig. 3a). The analysis highlighted the upregulation of tumor-associated macrophage markers and pro-tumoral markers in the CD74^+^ microglia/macrophages (Supplementary Fig. 3b-f, Table 2). In addition, differentially expressed genes in CD74^+^ microglia/macrophages (Fig. 2b, Table 3) were scored in a cohort of breast cancer brain metastases with clinical follow-up and matched primary tumors ^54^. This analysis identified a 37 gene-signature that stratifies patient survival from the diagnosis of the brain metastasis (Fig. 2c). In contrast, the clinical correlation was not reproduced when evaluated in the bulk RNA-seq of the matched breast cancer primary tumor (Fig. 2d), suggesting an organ-specific contribution of CD74^+^ microglia/macrophage.

**Figure 2.**
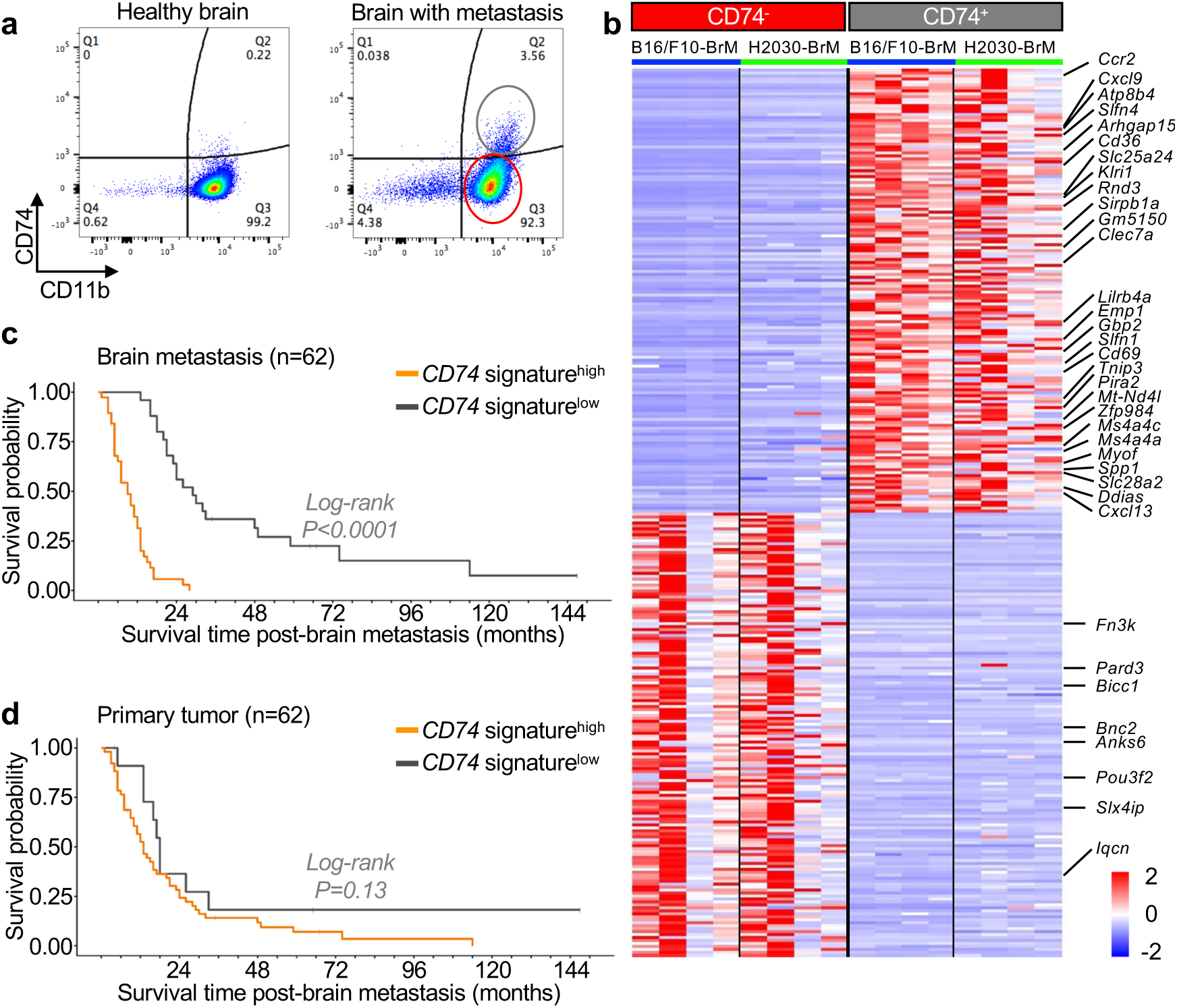
The pro-metastatic signature of reprogrammed CD74^+^ macrophages/microglia has clinical relevance. **a**. Representative flow cytometry plot from healthy and H2030-BrM metastatic brains. Each graph represents a pool of 3 brains. CD74^−^ microglia/macrophages are outlined in red, and CD74^+^ in gray. **b.** Heatmap of the top 150 upregulated and 150 downregulated genes from bulk RNA-seq analysis of CD74^+^ microglia/macrophages. Genes listed on the right constitute the 37-gene human signature. **c.** Survival graph of 62 patients with brain metastasis from breast cancer. CD74 signature was scored in RNA-seq data from brain metastasis human samples. The orange line represents patients enriched in the 37-gene signature derived from the CD74^+^ bulk RNA-seq signature (Fig. 2b), and the black line represents patients with lower enrichment of this signature. **d.** Survival graph of 62 patients with brain metastasis from breast cancer. CD74 signature was scored in RNA-seq data from breast primary tumors (matched samples from Fig. 2c). The orange line represents patients enriched in the 37-gene signature derived from the CD74+ bulk RNA-seq signature (Fig. 2b), and the black line represents patients with lower enrichment of this signature

Previously generated scRNA-seq data in brain metastasis ^20^ allowed us to interrogate the CD74^+^ microglia/macrophages. As expected, we identified the expression of *Cd74* distributed between microglia and bone marrow-derived macrophages (Supplementary Fig. 4a, b). Although some expression could also be detected in the astrocyte cluster (Supplementary Fig. 4b) as it has been previously reported ^55^, we could not confirm at the protein level the presence of CD74^+^ astrocytes surrounding brain metastasis (Supplementary Fig. 4c). We next separated these populations according to expression of microglia signature (Supplementary Fig. 4d) and bone-marrow derived macrophage signature (Supplementary Fig. 4e) previously described^56^. We observed that *Cd74* expression mainly occurred in the bone-marrow derived macrophage cluster (Supplementary Fig. 4f). We confirmed the predominance of bone marrow origin over microglia within the brain metastasis-associated CD74^+^ population in experimental models *in situ* using immunofluorescence and flow cytometry (Supplementary Fig. 4g-k).

To further dissect whether CD74^+^ bone marrow-derived macrophages alone, as the predominant CD74^+^ cell type within the metastasis associated microenvironment, reproduced the clinical correlation (Fig. 2c), we first separated BMDM from microglia by the expression of *Ccr2* ^57–59^ (Supplementary Fig. 4l). Subsequently, we identified differentially expressed genes (DEGs) between *Ccr2^+^; Cd74^+^* and *Ccr2^−^; Cd74^+^* populations (Table 4). Remarkably, a signature associated with *Ccr2^+^; Cd74^+^*, which we use as a strategy to score CD74^+^ bone marrow-derived macrophages, best recapitulated the clinical correlation in brain metastasis (Supplementary Fig. 4m), which also did not score when evaluated at the primary tumor (Supplementary Fig. 4n).

Thus, our data confirmed the clinical value of a signature derived from CD74^+^ microglia/macrophage and pointed to CD74^+^ bone marrow-derived macrophages as a contributor to the poor prognosis signature in brain metastasis.

### The protumoral activity of CD74^+^ macrophages/microglia relies on mitochondria-mediated plasticity

The transcriptomic analysis of CD74^+^ cells revealed a strong upregulation of the oxidative phosphorylation (OXPHOS) process (Supplementary Fig. 3a, Supplementary Fig. 5a, Table 5). This agrees with the primary use of this pathway as a source of energy in pro-tumoral macrophages ^60,61^. Due to CD74 overlap with *Ccr2* expression (Supplementary Fig. 4l) and Ccr2 protein levels (Supplementary Fig. 5b) and the observed clinical correlation with *Ccr2^+^; Cd74^+^* BMDM (Supplementary Fig. 4m), we decided to evaluate whether the MIF-induced acquisition of pro-metastatic functions could be reverted by genetically targeting components of the mitochondria required for OXPHOS mediated plasticity.

Beyond the enrichment of OXPHOS in the CD74^+^ microglia/macrophages bulk RNA-seq (Supplementary Fig. 3a, Supplementary Fig. 5a, Table 5), we also identified the Oma1-related pathway (Fig. 3a). Oma1 is a major determinant of mitochondria quality and critical to ensure its ability to sustain OXPHOS mediated cellular plasticity ^62^. To study the potential dependency of the pro-tumoral role of CD74^+^ macrophages on mitochondria mediated plasticity we performed brain metastasis *in vivo* assays using the Ccr2-Cre^ERT2^;Oma1^loxP/loxP^ genetically engineered mouse model (GEMM) (Fig. 3b). The GEMM showed a decreased viability of cancer cells in the brain as evaluated with both bioluminescence and histology (Fig. 3c, d and Supplementary Fig. 5c-e), however without affecting extracranial metastases (Supplementary Fig. 5f, g). Histological analysis demonstrated similar amount of brain metastasis-associated CD74^+^ microglia/macrophages compared to the wild-type mice (Fig. 3e, f), however a reduced presence of the NF-κB-reporter gene midkine (MDK) was detectable in CD74^+^ microglia/macrophages in the GEMM (Fig. 3g, h).

**Figure 3.**
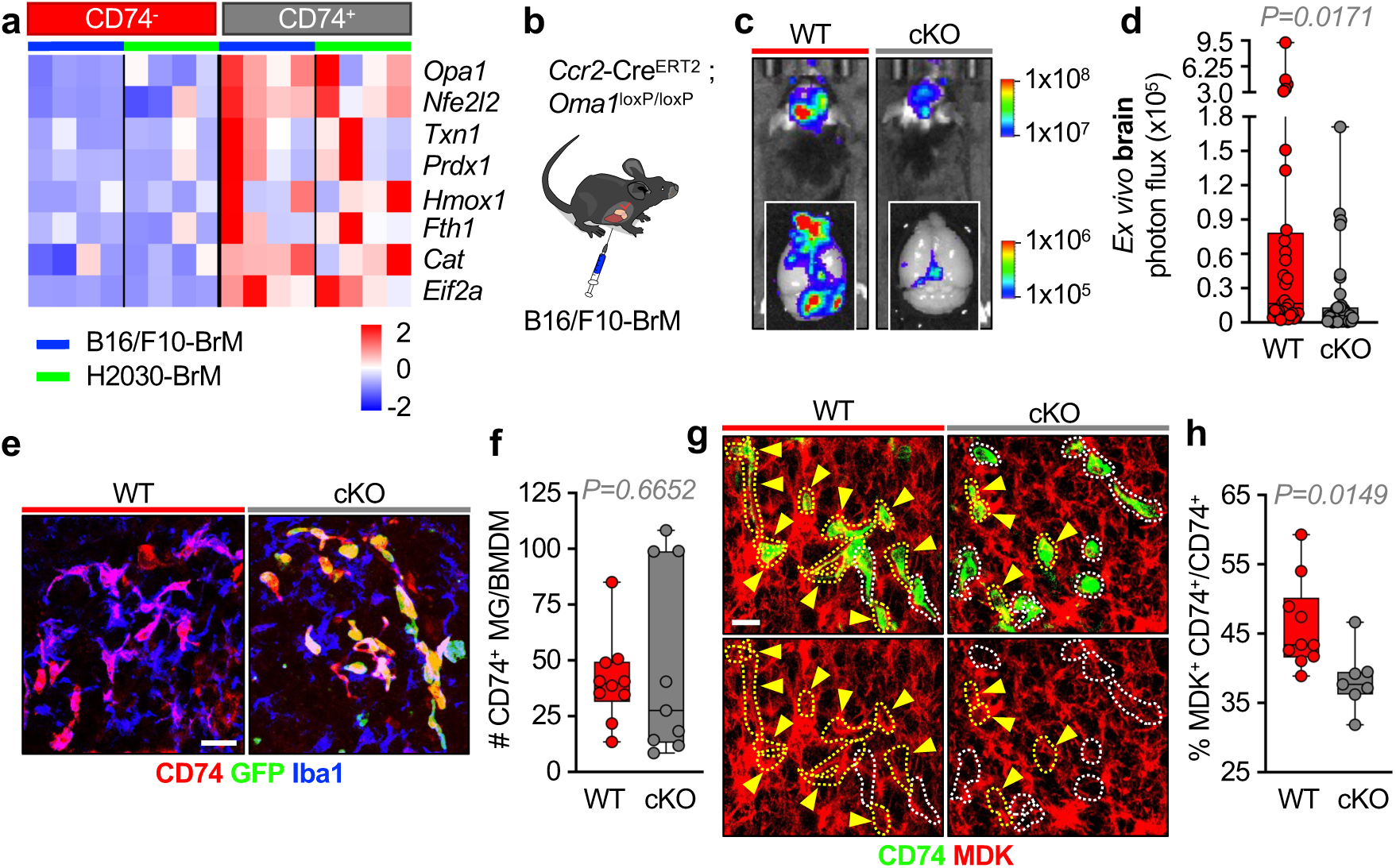
Brain metastasis-promoting activity depends on CD74^+^ macrophages/microglia mitochondria-mediated plasticity. **a**. Heatmap of Oma1 pathway-related genes that are upregulated in the CD74^+^ vs CD74^−^ microglia/macrophages. **b.** Schema of the mouse model. Mice carry a loxP site flanking the *Oma1* sequence, and Cre under the *Ccr2* promoter. Mice were intracardially injected with B16/F10-BrM cells and tamoxifen was administered to induce Cre expression 3 days after intracardiac injection. **c.** Representative BLI *in vivo* images of WT and cKO^CCR2^-*Oma1* mice 14 days after intracardiac injection. *Ex vivo* BLI of brains (down) are also shown. **d.** Quantification of *ex vivo* BLI of brains from c. Values are shown in box-and-whisker plot where each dot represents a different animal and the line corresponds to the median. n=33 (WT) and 35 (cKO) mice from 3 independent experiments. *P* value was calculated using an unpaired two-tailed t-test. **e.** Immunofluorescence against CD74, GFP (labeling Cre expression under the *Ccr2* promoter) and Iba1 in B16/F10-BrM tumors from WT and cKO mice. **f.** Quantification of images shown in e, representing the total numbers of CD74^+^ microglia/macrophages (MG/BMDM). Values are shown a in box-and-whisker plot where each dot represents the mean number of cells per brain, and the line corresponds to the median. n=10 (WT) and 9 (cKO) mice, 3-6 FOV per mouse, from 2 independent experiments. *P* value was calculated using an unpaired two-tailed t-test. **g.** Immunofluorescence against CD74 and midkine (MDK) in tumors. Dotted lines surround the CD74^+^ cells that are MDK^+^ (yellow) or MDK^−^(white). Double-positive CD74+ MDK+ are also indicated with a yellow arrow. Images with the red-channel only (MDK) are shown for clarity. Evaluation of the expression was done in Iba1^+^ cells. Scale bar: 10 µm. **h.** Quantification of images shown in g representing the percentage of CD74^+^ MDK^+^ microglia/macrophages (MG/BMDM) out of the total population of CD74^+^ Iba1^+^ cells. Values are shown in a box-and-whisker plot where each dot represents the mean percentage of double positive CD74^+^ MDK^+^ cells per brain, and the line corresponds to the median. n=10 (WT) and 9 (cKO) brains, 4-12 FOV per mouse, from 2 independent experiments. *P* value was calculated using an unpaired two-tailed t-test.

Thus, our data demonstrate that we can challenge brain metastasis viability by targeting mitochondria-mediated plasticity in CD74^+^ bone marrow-derived macrophages using a Ccr2-driven genetic strategy.

### Ibudilast challenges experimental brain metastases blocking CD74^+^ macrophages/microglia reprogramming

Given the potential to target a pro-metastatic component of the microenvironment of secondary brain tumors (Fig. 1b), which is absent from the healthy brain (Supplementary Fig. 1f), and the availability of a safe brain-penetrant compound ^63^ tested in other brain disorders (NCT03782415, NCT01982942), we decided to exploit the therapeutic potential of blocking the MIF-CD74 axis by using ibudilast. Ibudilast is a small molecule compound, which prevents binding of MIF to the its cognate receptor^43^. First, we confirmed that ibudilast added to the cancer cells *in vitro* did not compromise their viability (Supplementary Fig. 6a-c). In contrast, *in vivo* treatment in two experimental brain metastases models (H2030-BrM and B16/F10-BrM) with ibudilast reproduced the phenotype previously obtained by targeting MIF in cancer cells (Fig. 1e-g and Supplementary Fig. 2e, h-k) as measured by both bioluminescence (Fig. 4a, b and Supplementary Fig. 6d, e) and histology (Supplementary Fig. 6f-k). Similarly to the genetic approach targeting MIF in cancer cells, a modest reduction of extracranial metastasis was detected (Supplementary Fig. 6l-o).

**Figure 4.**
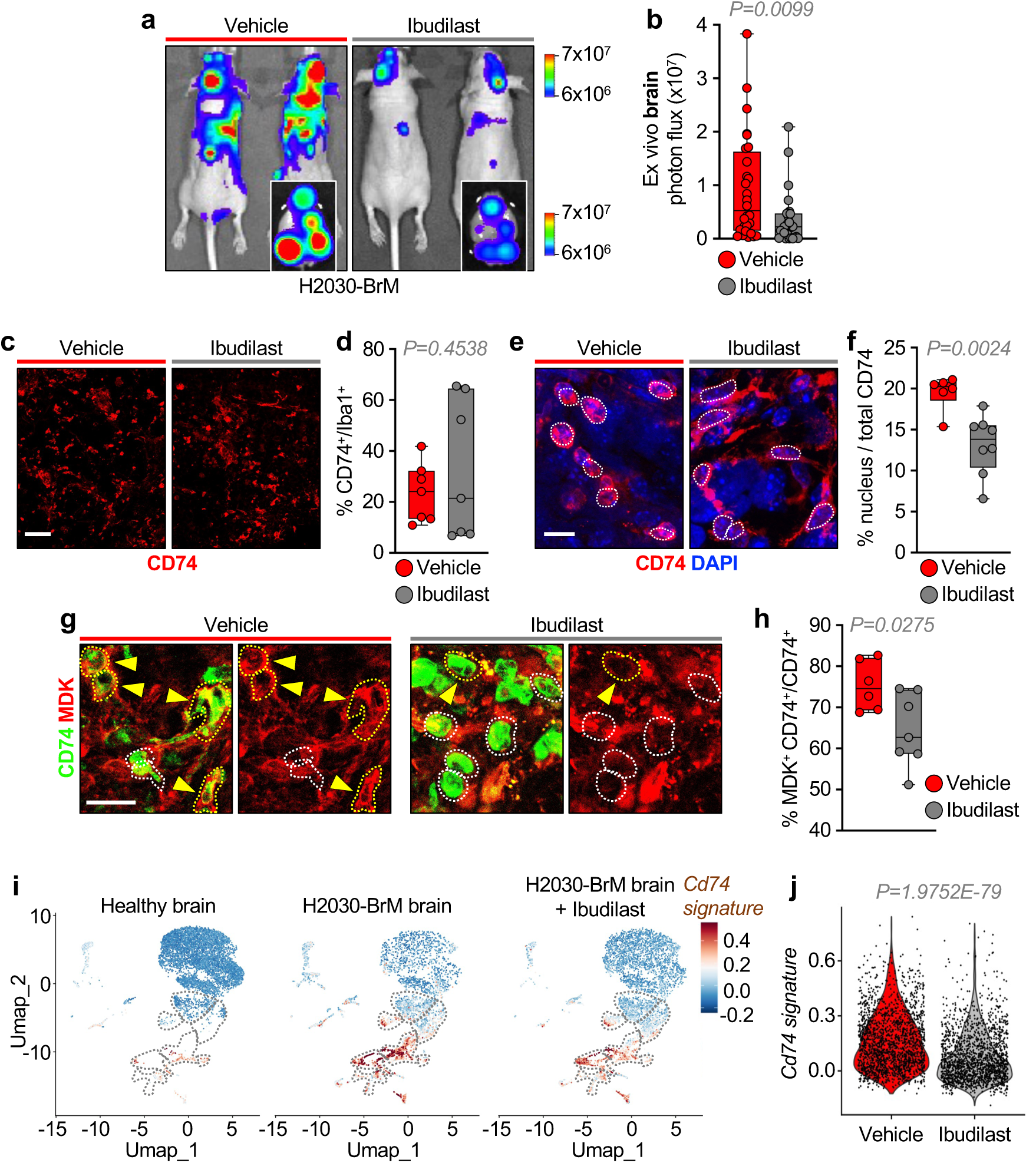
Ibudilast challenges brain metastases by targeting reprogrammed CD74^+^ macrophages/microglia. **a**. Representative BLI *in vivo* images of vehicle– or ibudilast-treated mice at endpoint (35 days after intracardiac injection). *Ex vivo* BLI of brains are also shown. **b.** Quantification of *ex vivo* BLI of brains from a. Values are shown in a box-and-whisker plot where each dot represents a different animal and the line corresponds to the median. n=28 (vehicle) and 27 (ibudilast) mice from 3 independent experiments. *P* value was calculated using an unpaired two-tailed t-test **c.** Immunofluorescence against CD74 in H2030-BrM tumors from vehicle– and ibudilast-treated mice. Scale bar: 75 µm. **d.** Quantification of images shown in c. representing the percentage of CD74^+^ area normalized to the Iba1^+^ area. Values are shown in a box-and-whisker plot where each dot represents the mean percentage of CD74^+^ normalized to the Iba1+ area per brain, and the line corresponds to the median. n=7 mice, 4-8 FOV per mouse. *P* value was calculated using an unpaired two-tailed t-test. **e.** Immunofluorescence images of CD74 and DAPI in H2030-BrM tumors from vehicle– and ibudilast-treated mice. White dotted lines outline the DAPI^+^ nuclei. Scale bar: 10 µm. **f.** Quantification of CD74-ICD found in the nucleus (DAPI) vs the membrane/cytoplasm on images shown in e. Measures were performed in Iba1^+^ cells by an automatic software. Values are shown in a box-and-whisker plot where each dot represents the mean percentage of CD74-ICD found in the nuclei per animal, and the line corresponds to the median. 10 FOV were captured per brain, in n=6 (vehicle) and n=8 (ibudilast) mice. *P* value was calculated using an unpaired two-tailed t-test of each experimental condition against control. **g.** Immunofluorescence against CD74 and MDK in Iba1^+^ cells in H2030-BrM tumors from vehicle– and ibudilast-treated mice. White dotted lines outline CD74^+^ MDK^−^ cells, while yellow dotted lines indicated with an arrow delineate the double positive CD74^+^ MDK^+^ cells. Images with the red-channel only (MDK) are shown for clarity. Evaluation of the expression was done in Iba1^+^ cells. Scale bar: 10 µm. **h.** Quantification of images shown in g, values are shown in a box-and-whisker plot where each dot represents the mean percentage of Iba1^+^ CD74^+^ cells positive for MDK normalized to the total CD74^+^ Iba1^+^ per brain. 12 FOV were captured per brain, in n=6 (vehicle) and n=7 (ibudilast). *P* value was calculated using an unpaired two-tailed t-test of each experimental condition against control. **i.** UMAP showing the expression of *CD74* signature obtained by bulk RNA sequencing (Fig. 2a, b and Supplementary Fig. 3a) in CD45^+^ cells found in the healthy and brain metastasis microenvironment of H2030-BrM model, categorized by condition: left: healthy brain, middle: H2030-BrM metastatic brain, and right: H2030-BrM metastatic brain treated with ibudilast. **j.** Violin plot presenting the distribution of *Cd74* signature score in the myeloid CD74^+^ populations (disease-associated microglia, reactive microglia and reactive macrophages) in H2030-BrM vehicle– and ibudilast-treated brains.

Importantly, ibudilast did not alter the amount of CD74^+^ microglia/macrophages *in vivo* (Fig. 4c, d and Supplementary Fig. 6p, q), but mimicked MIF knockdown phenotype by reducing both nuclear CD74-ICD levels in microglia/macrophages (Fig. 4e, f) and MDK expression, as a reporter of decreased NF-κB activity (Fig. 4g, h and Supplementary Fig. 6r, s).

To confirm that the reduction in brain metastasis burden observed *in vivo* was not mediated by ibudilast’s additional inhibition of phosphodiesterases (PDEs), we tested AV1013, an analog with significantly reduced PDE-inhibitory activity ^43^. AV1013 replicated ibudilast’s anti-metastatic effect in organotypic brain slice cultures (Supplementary Fig. 7a–c), discarding PDE inhibition as the primary driver of ibudilast’s action and rather reinforcing the therapeutic benefit linked to MIF blockade.

To further dissect the mechanism of action of ibudilast, we performed scRNA-seq of CD45^+^ sorted cells from the brain metastasis microenvironment (Supplementary Fig. 7d, e). Ibudilast-treated brains showed a clear shift in the immune landscape, with a nearly twofold increase in homeostatic microglia, and a marked decrease in reactive macrophages and infiltrative populations, showing a regression to control levels (Supplementary Fig. 7e, f). Notably, several subpopulations of microglia and macrophages in vehicle-treated mice expressed the protumorigenic signature identified in CD74^+^ microglia/macrophages (Fig. 2b, Fig. 4i), mimicking the emergence of a disease-associated cell type. The effect of ibudilast reducing the expression of the CD74 pro-metastatic signature was evident (Fig. 4i, j), which suggests that its anti-metastatic effect relies on its ability to impair MIF-dependent reprogramming of CD74^+^ macrophages/microglia (Fig. 1e-k, Supplementary Fig. 2a-c).

Furthermore, analysis of reactive macrophages, as the CD74^+^ cell subpopulation with the highest expression of the signature (Fig. 4i, Supplementary Fig. 7g), identified additional potential biomarkers of response (Supplementary Fig. 7h, Table 6). We validated *Cd274* (PD-L1) in CD74^+^ microglia/macrophages (Supplementary Fig. 7i, j) given its implications on local immunosuppression. Based on this finding, it is tempting to speculate that ibudilast would facilitate the anti-metastatic action of immunotherapies locally, at least in part, by limiting the expression of immune check-point blockade molecules.

Encouraged by this paradoxical finding on rewiring CD74^+^ microglia/macrophages from an antigen presenting cell into an immunosuppressive compartment of the metastasis microenvironment, we further investigated a potential correlation of ibudilast anti-metastatic phenotype on cytokine profiling associated with brain metastases (Supplementary Fig. 7d-i). Despite overall similar cytokine profiles between vehicle– and ibudilast-treated brains compared to naïve brains (Supplementary Fig. 8a), IL-2 stood out as the most dynamically altered. Its levels were markedly reduced in metastasis-bearing brains compared to healthy controls, and this drop was largely reversed by ibudilast treatment, restoring IL-2 to levels comparable to metastasis-naïve brains. While other cytokines showed similar trends, none matched the magnitude of IL-2’s decrease and subsequent recovery (Supplementary Fig. 8a). IL-2 plays a major role as an activator of multiple immune cells ^64^. Given our interest to understand the pro-metastatic roles of ibudilast both in immunocompetent but also in immunosuppressed models of brain metastases (Fig. 4a, b and Supplementary Fig. 6d, e) we explored the potential impact on natural killer (NK) cells. Both mouse and human NK cells activation and expansion are IL-2 dependent ^65,66^ and have been shown to influence brain metastasis progression ^67^. We assessed the numbers of NK cells in vehicle– and ibudilast-treated mice. We observed a significantly higher infiltration of NK cells in the Ibudilast-treated mice compared to the untreated controls (Supplementary Fig. 8b, c).

These results suggest that ibudilast inhibition of the MIF-CD74 signaling axis in microglia/macrophages impairs acquired pro-metastatic functions (MDK, PD-L1) but also suggests a broader influence in the brain metastatic niche toward a less supportive state for tumor growth (increased NK cells).

### Use of ibudilast on human brain metastases depicts a tumor agnostic strategy guided by biomarkers of therapeutic response

To evaluate the effect of ibudilast on human brain metastases, we used patient-derived organotypic cultures (PDOC) from freshly resected brain metastases obtained in RENACER ^68^.

Brain metastasis PDOC were established from 26 patients, capturing the clinical and biological heterogeneity of the disease across primary tumor types and clinical management (Table 7). The rationale for such tumor-agnostic approach derives from the broad and abundant presence of CD74^+^ microglia/macrophages among human brain metastases (Fig. 5a, Table 8) and our preclinical functional findings in both lung and melanoma brain metastases (Fig. 4a, b, Supplementary Fig. 6d, e). The 26 PDOC were tested for the sensitivity to ibudilast using a well-established pipeline ^69,70^ that consist on the incubation of the fresh tissue slice with the drug over a 3 days period (Fig 5b). Remarkably, 19/26 (73.08%) of the patient-derived organotypic cultures responded to treatment, defined as a reduction in cancer cell viability greater than 30%, following clinical benchmarks ^71^ (Fig. 5c, d).

**Figure 5.**
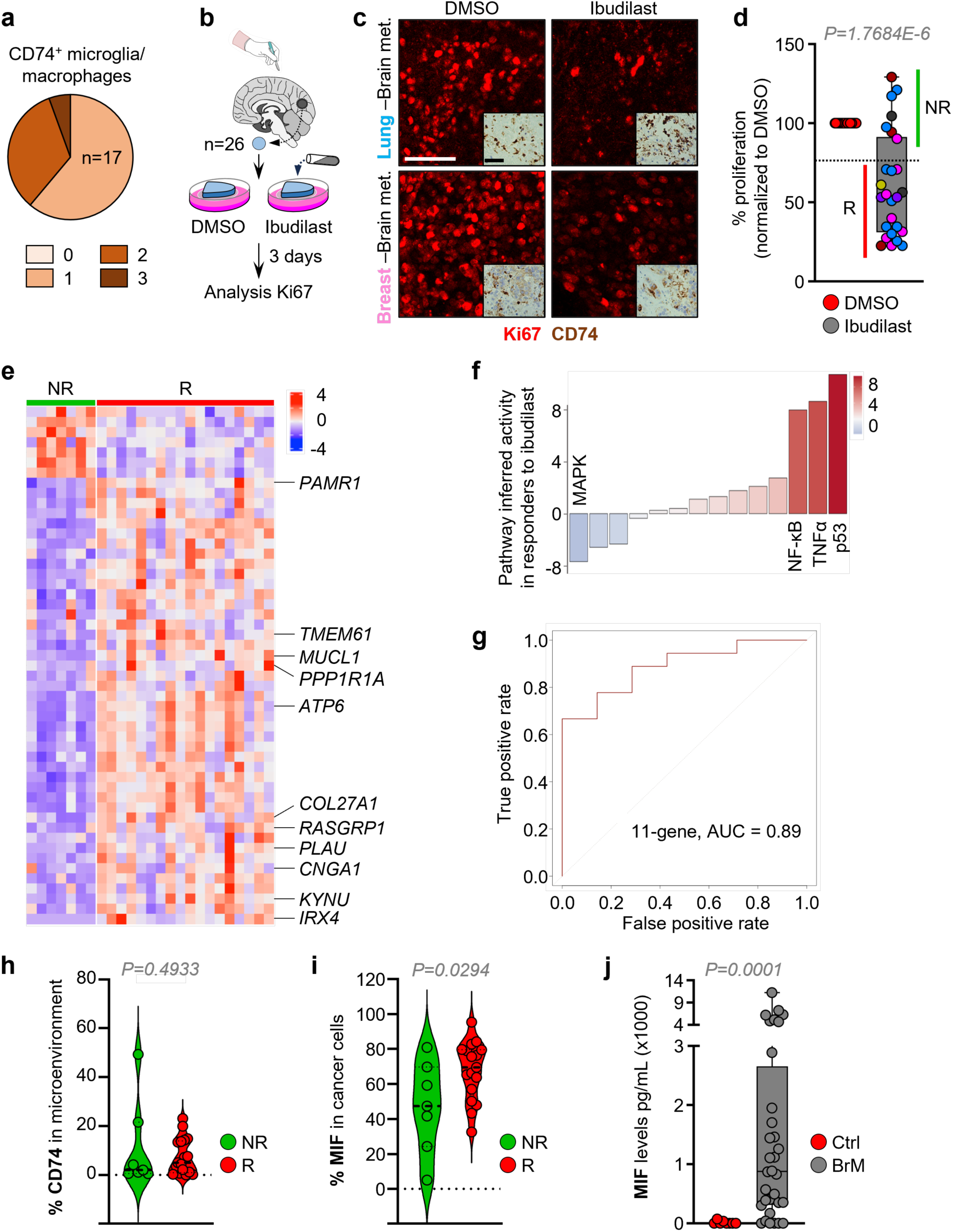
*Ex vivo* clinical use of ibudilast on patient-derived organotypic cultures suggests a tumor agnostic strategy against brain metastases. **a**. Quantification of CD74 in microglia/macrophages in human brain metastasis. 17/17 (100%) showed positive staining for CD74, from which 10/17 (58.82%) scored with 1, 6/17 (35.29%) with 2 and 1/17 (5.88%) with 3 according to the abundance and signal intensity of CD74 in the microenvironment. **b.** Schema of the experimental design. Fresh neurosurgeries from brain metastasis patients cut with a vibratome were cultured to generate patient-derived organotypic cultures (PDOC). The slices were cultured with ibudilast 50 μM for 3 days. **c.** Representative immunofluorescence against Ki67 labeling the proliferating cancer cells in PDOC derived from a lung cancer brain metastasis (top) and a breast cancer brain metastasis (bottom) after 3 days of *ex vivo* treatment with ibudilast. Scale bar: 75 μm. On the right down corner of each image immunohistochemistry staining against CD74 in the corresponding PDOC. Scale bar: 50 μm. **d.** Quantification of the relative number of Ki67^+^ cancer cells in PDOCs after treatment with ibudilast respect to the corresponding vehicle-treated PDOC with DMSO (from pictures shown in c). Dots are colored according to the primary source of the metastasis: blue: lung cancer. 42.31% (11/26), pink: breast cancer 26.92% (7/26), dark red: colorectal cancer 11.54% (3/26), purple: uterine cancer 7.69% (2/26), black: melanoma 7.69% (2/26), yellow: prostate cancer 3.85% (1/26). Values are shown in a box-and-whisker plot where each dot represents a different patient (mean value obtained from all PDOC from the same condition and patient) and the line in the box corresponds to the median. n= 26 patients, 3 slices per condition (DMSO or ibudilast) and 3 FOV per slice. *P* value was calculated using a paired two-tailed t-test. **e.** Heatmap of differentially expressed genes in non-responders (NR) vs ibudilast-responders (R) from samples in d, as determined by bulk RNA-seq performed on fresh, untreated neurosurgical samples. The 11 genes composing the ibudilast-response prediction signature are listed on the right. **f.** Pathway analysis of differential induced pathways in ibudilast-responders vs non-responders. **g.** AUC-ROC curve for the 11-gene signature, obtained by comparing non-responders vs. ibudilast responders, demonstrating its ability to predict treatment response. **h.** Quantification of CD74^+^ cells in the microenvironment in paraffin sections from ibudilast non-responders (n=7) and responders (n=17) patients. Values are shown in a violin plot where each dot represents one human sample, and the line corresponds to the median. *P* value was calculated using an unpaired two-tailed t-test. **i.** Quantification of MIF^+^ cells cancer cells in paraffin sections from ibudilast non-responders (n=7) and responders (n=17) patients. Values are shown in a violinplot where each dot represents one human sample, and the line corresponds to the median. *P* value was calculated using an unpaired two-tailed t-test. **j.** Measure of MIF levels (pg/mL) determined by ELISA in CSF from healthy donors and brain metastasis patients. Values are shown in a box-and-whisker plot where each dot represents a patient and the line corresponds to the median. *P* value was calculated using a Mann-Whitney U test.

Importantly, 53.85% of the organotypic cultures showed a reduction of 50% or more, with responses observed across multiple primary tumor origins (Fig. 5d). We then examined the 34.62% brain metastases that do not respond to ibudilast, classified as non-responders, and identified a transcriptomic signature that separates them from responders (Fig. 5e, f). Interestingly, a reduced panel of just 11 genes was sufficient to predict response, offering a clinically actionable biomarker set (Fig. 5e, g).

The potential to develop a clinical trial using ibudilast in brain metastasis based on its brain penetrance, safety profile ^44,47,63^ and its potential to facilitate immunotherapies (Supplementary Fig. 7h–j), encouraged us to identify additional biomarkers to select patients. We evaluated whether protein levels of CD74 or MIF correlated with ibudilast response. While CD74 expression levels did not significantly differ between responders and non-responders (Fig. 5h), MIF levels were significantly higher in responders (Fig. 5i), suggesting that MIF abundance might be a predictive marker of treatment efficacy. However, relying on tumor tissue biopsy for biomarker detection requires invasive procedures. Alternatively, the secreted nature of MIF (Supplementary Fig. 1d) and its high levels in brain metastatic cells (Fig. 1a and Supplementary Fig. 1a-e), suggested its potential as a biomarker compatible with liquid biopsy. CSF from non-tumor hosting patients (intracerebral pressure diagnosis, Table 9) and from RENACER patients (Table 9) were evaluated with a MIF specific ELISA. Remarkably, MIF levels were increased only in patients with brain metastases (Fig. 5j), confirming its potential clinical value as a non-invasive liquid biopsy biomarker.

### CD74^+^ in macrophages/microglia is a biomarker of several brain disorders

Given that CD74^+^ microglia/macrophages have been broadly reported in other brain disorders beyond oncology ^3–6^, and generally considered a marker of the MHC-II complex in spite of the limited functional evaluation for its contribution to the disease, we asked whether our findings on brain metastasis-associated CD74^+^ microglia/macrophages could be applied to this cell subpopulation present in Alzheimer’s disease (AD) and multiple sclerosis (MS).

First, we confirmed that the experimental models of AD and MS did show CD74^+^ microglia/macrophages in areas affected by the pathological findings (i.e., phospho-Tau and myelin debris, respectively) (Fig. 6a-c) as well as MIF-positive areas (Fig. 6d-f). Furthermore, our comparative scRNA-seq approach on CD45^+^ cells (Supplementary Fig. 9a-f) allowed us to demonstrate an expansion of *Cd74* expression in all disease models compared to their aged-matched controls (Fig. 6g-l).

**Figure 6.**
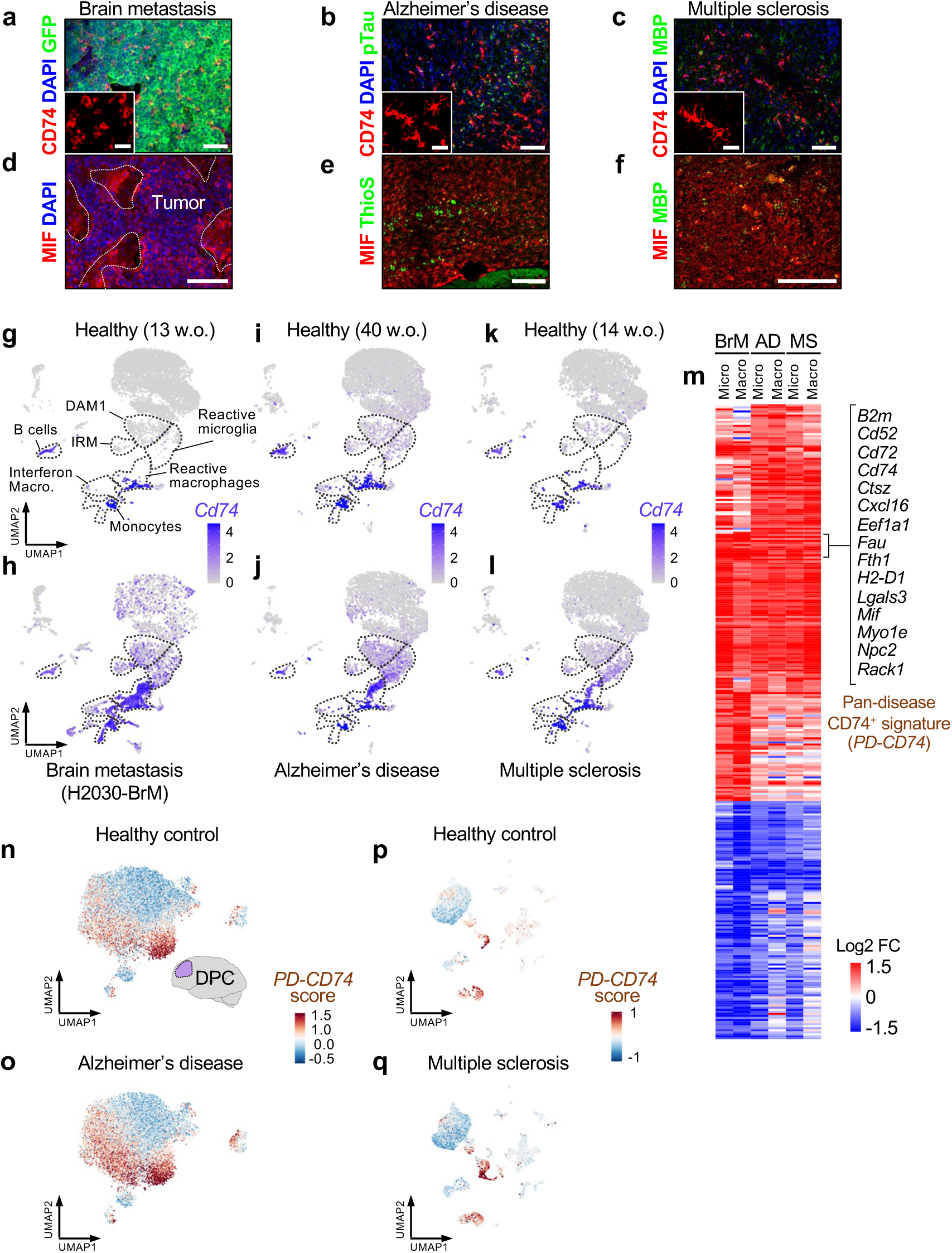
CD74^+^ in macrophages/microglia is a biomarker of brain disorders. **a**. Immunofluorescence against CD74, GFP (H2030-BrM cells) and DAPI in H2030-BrM metastatic brain. Scale bar: 75 µM. Magnification scale bar: 25 µM. **b.** Immunofluorescence against CD74, pTau and DAPI in brain from an Alzheimer’s disease mouse model. Scale bar: 75 µM. Magnification scale bar: 25 µM. **c.** Immunofluorescence against CD74, MBP (myelin basic protein) and DAPI in a brain from a cuprizone murine model of MS. Scale bar: 75 µM. Magnification scale bar: 25 µM. **d.** Immunofluorescence against MIF and DAPI in H2030-BrM metastatic brain. White dotted lines outline the tumoral area. Scale bar: 25 µM. **e.** Immunofluorescence against MIF and Thioflavin S (ThioS) in brain (somatosensory cortex) from an Alzheimer’s disease mouse model. Scale bar: 25 µM. **f.** Immunofluorescence against MIF and myelin (MBP) in a demyelinating plaque from a secondary progressive MS patient. Scale bar: 25 µM. **g. – l.** *Cd74* expression in UMAPs of CD45^+^ populations found in healthy brain from immunocompromised mice (**g**), in H2030-BrM metastatic brains (**h**), in healthy aged brain from immunocompetent mice (**i**), in the brains of mouse models of Alzheimer’s disease (**j**), in healthy brain from immunocompetent mice (**k**), and in the brains of mouse models of multiple sclerosis (**l**). DAM1: disease-associated microglia. Interferon Macro.: interferon-responsive macrophages. **m.** Heatmap depicting differentially expressed genes with significant upregulation (adjusted p-value < 0.05, log2 fold change > 0.5) and significant downregulation (adjusted p-value < 0.05, log2 fold change < –0.5) shared by microglia and macrophages across each disease context (BrM: brain metastasis, AD: Alzheimer’s disease, MS: multiple sclerosis). The common signature of 15 protein-coding genes exhibiting significant differential enrichment (adjusted p-value < 0.05, log2 fold change > 0.5) across all contrasts is label as the pan-disease CD74+ gene signature (PD-CD74).. **n. – q.** UMAPs showing the cell scores for the experimental pan-disease CD74 gene signature (*PD-CD74*) from m. in human brain scRNA-seq data. **n.** from the dorsolateral prefrontal cortex (DPC) of healthy individuals, **o.** from the DPC of Alzheimer’s disease patients, **p.** from healthy individuals, **q.** from patients with multiple sclerosis,

Analysis of upregulated pathways in the CD74^+^ versus CD74^−^ myeloid compartments revealed signatures that are also part of pro-tumorigenic macrophages in tumors such as NF-κB ^72^, mTOR ^73^ or oxidative phosphorylation ^60,61^ (Table 10).

Importantly, gene expression analysis in CD74^+^ versus CD74^−^ microglia/macrophages suggested the existence of a conserved 67-gene signature across disorders, including 15 protein-coding genes (Fig. 6m) (Table 11). We refer to this shared program composed of 15 genes as the pan-disease CD74^+^ signature (PD-CD74). The experimental pan-disease CD74^+^ signature was evaluated in publicly available datasets of patients affected with AD ^74^ and MS ^75^. Remarkably, the PD-CD74 proved to be enriched in human brains affected with AD/MS in comparison with healthy individuals (Fig. 6n-q, Supplementary Fig. 9g-h).

Our data suggest that CD74^+^ microglia/macrophages represent a subpopulation of disease-associated cells enriched in a variety of brain disorders that potentially contribute to the human disease.

## Discussion

Our findings demonstrate the pro-metastatic nature of CD74^+^ microglia/macrophages, challenging the prevailing assumption of being a surrogate of MHC-II activity ^3–6^. While CD74 is a known MHC-II chaperone, it can also act as a cytokine receptor for MIF. Upon ligand binding, CD74 is cleaved, and its intracellular domain (ICD) translocates to the nucleus, triggering a non-canonical transcriptional program via NF-κB ^24,52^. We confirmed this mechanism in CD74⁺ microglia/macrophages in brain metastases, using both nuclear staining and MDK expression as functional readouts of NF-κB activation.

Although MIF has previously been suggested to originate from the brain metastatic niche ^19^, we provide functional evidence that metastatic cancer cells are the primary source of MIF in both experimental and patient-derived models. Importantly, this tumor-derived MIF rewires microglia/macrophages into a disease-promoting state. We also show that this switch depends on mitochondrial plasticity, as we can suppress nuclear translocation of the CD74-ICD-dependent NF-κB program upon targeting components that regulate mitochondrial quality.

Based on our single-cell data and depletion experiments, this phenotype appears to be predominantly mediated by bone marrow-derived macrophages; however, we cannot exclude a contribution from microglia. This is particularly relevant given recent findings in melanoma brain metastases showing that activation of the Rela/NF-κB pathway in microglia drives a pro-tumoral program, whereas its inhibition reprograms them toward a proinflammatory, anti-tumor state, enhancing responses to immune checkpoint blockade ^76^.

Although MIF–CD74 signaling has been implicated in tumorigenesis across multiple cancers, most studies outside of primary brain tumors have focused on CD74 expression in tumor cells themselves ^30,31,34,35^, rather than in the immune microenvironment. In contrast, gliomas have revealed roles for CD74 in glioma-associated microglia/macrophages (GAMs), with evidence for both immune suppression and tumor-promoting activity ^39,40^. Our work extends this concept beyond gliomas to brain metastases, where we identify a distinct, MIF-responsive CD74⁺ subpopulation of myeloid cells that drives disease progression. Moreover, while CD74–NF-κB activation may represent a general mechanism, our findings suggest that its pathological relevance is context-dependent. Specifically, we observed only modest effects in lung metastases, in strong contrast to the brain-specific phenotype using the different approaches. This suggests that the MIF–CD74 axis plays a particularly prominent role in the brain metastatic niche. Nevertheless, we cannot exclude its relevance in other metastatic sites or primary tumors without further investigation. Comparative studies across tumor types and compartments will be essential to delineate the spatial and functional specificity of this pathway.

In addition to the benefit provided by genetic targeting of MIF in cancer cells, we demonstrate the potential of a pharmacologic intervention strategy by using ibudilast, a brain-penetrant MIF-CD74 inhibitor already in clinical trials for glioblastoma. While ibudilast is known to sensitize glioblastoma cells to temozolomide and improve survival in preclinical models ^46^, the mechanism has remained unclear. In glioblastoma, ibudilast has been shown to reduce the immunosuppressive function of M-MDSCs and enhance CD8⁺ T cell activity in the tumor microenvironment ^77^. Moreover, additional studies using CD74-neutralizing antibodies or MIF siRNA in glioma have linked the MIF-CD74 pathway to tumor immune evasion and myeloid reprogramming ^40^, supporting the broader therapeutic relevance of MIF-CD74 signaling. Our findings point out to a broader influence of ibudilast on the immune landscape as depicted by its modulation on IL-2 levels, which correlated with increased NK cells infiltration.

To assess clinical relevance, we tested ibudilast in 26 PDOC from a range of primary tumors. Although most patients responded to ibudilast, the profiling of the metastases suggests a predictive molecular classifier based on 11 genes. Given the safety, brain penetrance and the molecular classifier of response together with the liquid biopsy biomarker that could facilitate patient selection and response follow up, we envision the potential of developing a clinical trial with ibudilast in patients with brain metastases.

Interestingly, ibudilast has also shown clinical benefit in progressive multiple sclerosis, slowing brain atrophy in Phase II trials ^44,47^. While the exact mechanism that explain the clinical benefit remains to be clarified, it is often assumed to be primarily mediated through ibudilast’s inhibition of phosphodiesterase (PDE). However, our findings raise the possibility that ibudilast’s benefit in MS may also involve modulation of disease-associated CD74⁺ microglia/macrophages. This same subpopulation, which we identify in both MS and Alzheimer’s disease models, shares a conserved NF-κB– driven signature that parallels the pro-metastatic program observed in brain metastases.

Taken together, our work identifies a pan-disease, reprogrammable subpopulation of CD74⁺ myeloid cells that drives pathology in brain metastases and potentially other neurological diseases. Therapeutically targeting this subpopulation via the MIF–CD74 axis could open new avenues in neuro-oncology, neurodegenerative and neuroinflammatory disorders. Comparative multi-disease analyses will be critical to validate these findings and broaden their clinical impact.

## Methods

### Animal studies

All animal experiments were performed in accordance with protocols approved by the CNIO, Instituto de Salud Carlos III and Comunidad de Madrid Institutional Animal Care and Use Committee (PROEX299/15 and PROEX135/19).

CX3CR1-EGFP mice were obtained from Jackson Laboratory (B6.129P-Cx3cr1tm1Litt/J; 005582). cKO-Oma1 were generated by breeding Ccr2-Cre/ERT2-GFP (C57BL/6-Ccr2 < em1(cre/ERT2)Peng> /J Stock No. 035229, Jackson Laboratory) with *Oma1^l^*^oxP/loxP^, kindly shared to Dr. José Antonio Enriquez (CNIC) by Dr. Thomas Langer (MPIBA). Alzheimer’s disease mice (P301S Tauopathy, C57BL/6 background) were provided by Dr. Jesús Avila’s lab (CBMSO). These transgenic mice develop neurological symptoms at 3 months, with a median lifespan of 9–12 months. The cuprizone demyelinating mice as MS model were provided by Dr. Fernando de Castro’s lab (Instituto Cajal-CSIC), as previously described ^78,79^ in agreement with protocols approved by the Instituto Cajal-CSIC, Consejo Superior de Investigaciones Científicas-CSIC and Comunidad de Madrid Institutional Animal Care and Use Committee (PROEX044_19 and PROEX030.1-24), demyelination was induced in 6-week-old male C57BL/6 mice by feeding them 0.2% cuprizone ad libitum for 5 weeks. Brain colonization assays were performed in C57BL/6 mice and athymic nu/nu (ENVIGO #069) 6–10 weeks as previously described ^19^, by injecting 100 μL of PBS into the left ventricle containing 100,000 cancer cells or 1 μL of PBS intracranially (right frontal cortex, approximately 1.5 mm lateral and 1 mm caudal from the bregma, and to a depth of 2 mm) containing 40,000 cancer cells by using a gas-tight Hamilton syringe and a stereotactic apparatus.

Metastatic colonization was analyzed *in vivo* weekly and *ex vivo* at the endpoint by BLI. Anesthetized mice with isoflurane were injected with D-luciferin (150 mg/kg) retro– orbitally and imaged in an IVIS machine. Average radiance (photons/sec/cm^2^/steradian) from each region of interest (ROI) was measured using Living Image software, version 4.5. ROIs were created in the 2D bioluminescent image in the head or body of each mouse for *in vivo* analysis or around the corresponding organ for ex vivo analysis. For intracranial injections, *ex vivo* values at the end point were normalized to the BLI values of the head *in vivo* 3 days after injection of the cancer cells.

Cre expression was induced by tamoxifen (TMX) 3 days after intracardiac inoculation of cancer cells, when it is considered that the first brain metastatic cells have crossed the blood brain barrier. TMX (T5648, Sigma-Aldrich) was prepared as a 20 mg/mL stock solution in corn oil. Mice were administered tamoxifen via intraperitoneal (i.p.) injection at a dose of 50 mg/kg body weight. The administration schedule consisted of 5 consecutive days of tamoxifen injection, followed by a 2-day rest period, and then an additional 5 days of tamoxifen injection, until the experimental endpoint. Ibudilast (S4837, Selleck Chemicals and CAS 50847-11-5, ChemSpace) was administered by oral gavage daily (20 mg/kg for immunocompromised models or 30 mg/kg for immunocompetent models), beginning 7 days (immunocompromised model) or 3 days (immunocompetent model) after intracardiac injection and treatment continued until mice reached the end point of the experiment.

For bone marrow transplant experiments host mice were sub-lethally irradiated with two 4.5-Gy doses within a 4-hour period. Bone marrow (BM) cells were harvested from donor mice by flushing the tibias and femurs with phosphate-buffered saline (PBS). The collected BM cells were then filtered through a 70 μm cell strainer. The required number of BM cells were resuspended in PBS for injection. For transplantation, the irradiated host mice were anesthetized, and BM cells were injected retro-orbitally in a volume of 100 μL PBS containing 2×106 cells. Three weeks after, B16/F10-BrM cells were intracardially injected to the mice.

### Mouse organotypic cultures

Organotypic cultures from adult mouse brains were prepared as previously described ^69^. In brief, brains were dissected in HBSS supplemented with HEPES (2.5 mM, pH 7.4), CaCl2 (1 mM), NaHCO3 (4 mM), MgCl2 (1mM) and D-glucose (30mM) embedded in 4% low-melting agarose (Lonza) preheated at 42°C. The embedded brains were cut into 250 mm slices using a vibratome (Leica) and each slice was divided in two pieces at the hemisphere. The slices were placed with flat spatulas on top of 0.8-μm pore membranes (Sigma-Aldrich) floating on slice culture media (DMEM, supplemented HBSS 27%, FBS 5%, D-Glucose (27 mM), L-glutamine (1 mM), and 100 IU/mL penicillin/streptomycin) and subjected to the corresponding procedure. For the IFN-γ experiments, slices derived from wild-type brains from C57BL/6 mice 6-8 weeks of age. Slices were cultured in the presence or absence of recombinant murine IFN-γ 250 ng/mL (315-05-100UG, PeproTech) and recombinant mouse MIF protein 100 ng/mL (1978-MF-0257CF, R&D Systems) for 12 hours. For some experimental conditions, bone marrow-derived cells were isolated from tibias and femur from wild-type CX3CR1-EGFP mice and place on top of the brain slice (5,000 cells suspended in 2 μL of RPMI medium).

When the experiments were finished, brain slices were fixed with 4% paraformaldehyde (PFA) overnight at 4°C and immunofluorescence was performed.

### Cell Culture

Brain metastatic cell lines (BrM) from human and mouse origins have been previously described. H2030-BrM3 (abbreviated as H2030-BrM) ^51^ cells were cultured in RPMI1640 media supplemented with 10% FBS, 100 IU/mL penicillin/streptomycin, 1 mg/mL amphotericin B and 2 mM L-glutamine. B16/F10-BrM7 (abbreviated as B16/F10-BrM) ^19^ cells were cultured in DMEM media supplemented with 10% FBS, 100 IU/mL penicillin/streptomycin, 1 mg/mL amphotericin B and 2 mM L-glutamine. HEK293T were cultured in DMEM media supplemented with 10% FBS, 100 IU/mL penicillin/streptomycin, 1 mg/mL amphotericin B and 2 mM L-glutamine. All the cell lines harbored a plasmid expressing luciferase under the PGK promoter.

For the *in vitro* proliferation assay, 1,000 B16/F10-BrM cells per well harboring either shControl (shCtrl) or sh*Mif*#1/#2 were seeded in black 96 well-plates (Fisher Scientific, ref. 08-772-225). Plates were scanned by the Opera Phenix Plus High-Content Screening (HCS) System (Perking Elmer) 12 hours, 5 days and 7 days after seeding. Cells were automatically counted using Harmony v.5.2 software.

B16/F10-BrM and H2030-BrM cells harboring MID KD were generated by lentiviral transduction. HEK293T cells at 70% confluence were transfected in Opti-MEM with Lipofectamine 2000 (Invitrogen) and incubated at 37°C overnight with 8.75 µg of the following plasmids: pMDLg/pRRE (#12251, Addgene), pRSV-Rev (#12253, Addgene), VSV.G (#14888, Addgene) and lentiviral vectors carrying the corresponding shRNA against human *MIF* (TRC21254, Horizon Discovery), mouse *Mif* (TRC142338, Horizon Discovery) or the corresponding non-targeting control. The following day, media was replaced with DMEM supplemented with 10% FBS and 2 mM L-glutamine and virus production was maintained for 36-48 h. Viral supernatant was collected, passed through a 0.45 µm syringe filter and added to H2030-BrM (human *MIF*) at 50% confluence in RPMI1640 supplemented with 10% FBS, 2 mM L-glutamine and polybrene (5 µg/mL; Sigma-Aldrich) or B16/F10-BrM (mouse *Mif*) at 50% confluence in RPMI1640 supplemented with 10% FBS, 2 mM L-glutamine and polybrene (5 µg/mL; Sigma-Aldrich). The following day, media was replaced with complete culture media. Selection with puromycin (2 µg/mL; Sigma-Aldrich) was started 24-48 h after and maintained until complete cell death was observed in the non-infected cancer cells.

### Immunofluorescence and immunohistochemistry

Tissue for free-floating immunofluorescence was processed after overnight fixation at 4°C with 4% paraformaldehyde (PFA). Slices for PDOCs were obtained using a vibratome (250 mm). Mice brains were processed, and mouse slices were performed by using a sliding microtome (80 mm, Thermo Fisher Scientific) or vibratome (250 mm). Slices were blocked using a blocking solution composed by 10% NGS, 2% BSA and a 0.25% Triton X-100 in PBS for 2 h at RT. Primary antibodies were incubated at 4°C overnight in the blocking solution and the following day 30 min at RT. The appropriate fluorochrome-conjugated secondary antibody was added in blocking solution and incubated for 2 h at RT after extensive washing with PBS-Triton 0.25%. After extensive washing with PBS-Triton 0.25%, using DAPI (1 mg/mL, Sigma-Aldrich) for 7 min at RT nuclei were stained. After extensive washing with PBS, slices were mounted on glass slides using Fluoromount-G mounting medium (SouthernBiotech).

| List of primary antibodies |  |  |  |  |
| --- | --- | --- | --- | --- |
| Antibody | IF dilution | IHC dilution | Provider | Reference |
| GFP | 1:1000 | - | Aves Labs | GFP-1020 |
| CD74 (mouse) | 1:500 | 1:2000 | BD Pharmigen | 555317 |
| CD74 (human) | 1:300 | 1:300 | Abcam | ab213104 |
| Iba1 | Mouse | 1:500 | WAKO | 019-19741 |
|  | Human | 1:300 |  |  |
| Ki67 | 1:500 | - | Abcam | ab15580 |
| HMB45 | 1:500 | - | Abcam | ab732 |
| Luciferase | 1:500 | - | Novusbio | NB100-1677 |
| MDK | 1:500 | - | CNIO | - |
| MIF | 1:100 | 1:100 | Santa Cruz | sc-271631 |
| GFAP | 1:1000 | - | Millipore | MAB360 |
| NK1.1 | 1:300 | - | Thermo Fisher | MA1-70100 |
| CD206 | 1:100 | - | BioRad | MCA2235T |
| PPARγ | 1:100 | - | Cell Signaling | 2435 |
| PD-L1 | 1:100 | - | Abcam | ab213480 |
| pTau<br>(pSer202+Thr205) | 1:1000 | - | Pierce Endogen | AT8 |
| MBP | 1:500 | - | Abcam | ab40390 |
| ThioS | 1:300 |  | Sigma | T1892 |

| List of secondary antibodies |  |  |  |
| --- | --- | --- | --- |
| Antibody | IF dilution | Provider | Reference |
| Anti-chicken Alexa-Fluor 488 | 1:500 | Invitrogen | A11039 |
| Anti-rabbit Alexa-Fluor 488 | 1:300 | Invitrogen | A32731 |
| Anti-rat Alexa-Fluor 488 | 1:300 | Invitrogen | A11006 |
| Anti-rabbit Alexa-fluor 555 | 1:300 | Invitrogen | A21428 |
| Anti-rat Alexa-Fluor 555 | 1:300 | Invitrogen | A21434 |
| Anti-mouse Alexa-Fluor 555 | 1:300 | Invitrogen | A21422 |
| Anti-rabbit Alexa-fluor 633 | 1:300 | Invitrogen | A21070 |
| Anti-mouse Alexa-Fluor 633 | 1:300 | Invitrogen | A21052 |
| Anti-mouse Alexa-Fluor 700 | 1:300 | Invitrogen | A21036 |
| Anti-rat Alexa-Fluor 488 | 1:300 | Invitrogen | A21208 |
| Anti-goat Alexa-Fluor 555 | 1:300 | Invitrogen | A21432 |

For immunofluorescence in paraffin sections, wild-type C57BL/6 mouse brain, liver, lymph node, pancreas, kidney, lung, intestine and spleen were formalin-fixed and paraffin-embedded. In both cases, sections (3 μm) were obtained. After deparaffinization and rehydration, antigen retrieval was performed by boiling in citrate buffer pH 6 for 10 min. Sections were incubated with 3% BSA / 0.1% Triton in PBS for 45 min at room temperature and incubated with primary antibodies overnight at 4°C. After washing with 0.1% Triton/PBS, the appropriate fluorochrome-conjugated secondary antibodies were added for 45min, sections were washed, and nuclei were counterstained with DAPI. After washing with PBS, sections were mounted with Prolong Gold Antifade Reagent (Life Technologies).

Immunohistochemistry of paraffin embedded tissues was performed at the CNIO Histopathology Core Facility. For the different staining methods, the slides were deparaffinized in xylene and rehydrated by a graded ethanol series to water. Several immunohistochemical reactions were performed on an automated immunostaining platform (Autostainer Link 48, Agilent; Discovery XT-ULTRA, Roche/Ventana). First, antigen retrieval was performed with the appropriate pH buffer and endogenous peroxidase was blocked (3% hydrogen peroxide). The slides were then incubated with the appropriate primary antibody for single or double staining. Following the primary antibody, the slides were incubated with appropriate secondary antibodies and with horseradish peroxidase (HRP)–conjugated visualization systems when needed.

The immunohistochemical reaction was revealed using ChromoMap DAB, DISCOVERY Purple, or Teal Kit (Roche/Ventana). Nuclei were counterstained with hematoxylin. Finally, slides were dehydrated, rinsed, and mounted for microscopic evaluation. Positive controls for primary antibodies were included in each staining series.

Lysozyme IHC and RNAscope staining method were performed in an automated immunostaining platform (Ventana DISCOVERY ULTRA, Roche), including deparaffinization and rehydration as a part of the platform protocol with the appropriate probe: probe against *Mif/MIF* mRNA (R&D systems, 513809 Q-284534 for mouse and 451049 Q-284534 for human). After the probe, slides were incubated with the corresponding probe amplification kit (RNAscope VS Universal HRP Reagent Kit, ACD, cat.# 323210), conjugated with HRP, and the reaction was developed using 3,3-diaminobenzidine tetrahydrochloride (DAB Detection Kit, Roche/Ventana, cat.# 760-224).

### Image acquisition and analysis

Immunofluorescence images were acquired using a Thunder Imaging System (Leica-Microsystems) equipped with AFC, LED8 excitation light source and a DFC9000GTC camera using 5x/NA 0.12 and 10x/NA 0.45 dry objective for whole slide imaging under the Navigator software integrated in the LAS X v 3.8.1. For z-stack confocal imaging a TCS SP8 STED 3X (Leica-Microsystems) equipped with AFC, a, a white light laser, and PMT and HyD SMD detectors under the Navigator software and a TCS SP5 (Leica-Microsystems) equipped, AOBS under the LAS AF v2.6 software were used with 20x/NA 0.7 dry, 20x/NA 0.75 Multiple and 63x/NA 1.4 Oil immersion objectives. For analyzing the CD74 nuclei translocation (Supplementary 2C), confocal microscopy images were analyzed in 2D using ImageJ and custom Java scripts. Nuclei and cytoplasm were segmented using Otsu thresholding and median filtering, followed by binary morphological operations to refine the segmented areas using MorphoLibJ library ^80^. Fluorescence intensity was quantified for each z-slice, with mean and sum values calculated for both nuclei and cytoplasm areas. GitHub repository at https://github.com/cnio-cmu-BioimageAnalysis/fluorescenceQuantification_code.

For detecting the different microglia positive populations, confocal microscopy images were analyzed using a combination of Groovy and Python scripts. Initial segmentation of cells was performed in 3D using the Deep Learning-based Cellpose ^81^ library with the cyto3 model to accurately identify microglia based on the Iba1 marker. The segmented labels were then processed and analyzed using the mcib3d ^82^ library. For each marker described, the mean and standard deviation of intensity values were calculated across the entire dataset of microglia masks in 3D. Cells were identified as positive for a given marker if their intensity exceeded the threshold of mean plus one standard deviation (approximately the 84th percentile). GitHub repository: https://github.com/cnio-cmu-BioimageAnalysis/3DmicrogliaAnalysis_code.

Immunohistochemistry images were captured with Zen Blue Software v3.1 (Zeiss). DAB immunohistochemistry intensity was measured using QuPath v.0.4.2.

### Processing of mouse brains for flow cytometry analyses

Mouse brains were extracted in pre-cooled D-PBS 1X and were processed with the Adult Brain Dissociation Kit (Miltenyi, ref. 130-107-677) using gentleMACS C Tubes (Miltenyi, ref. 130-093-237) and the gentleMACS™ Octo Dissociator (Miltenyi, ref. 130-096-427). The resulting homogenates were processed as previously described ^83^. Homogenates were filtered with a 70 μm strainer and were centrifuged at 300 g for 10 min at 4 °C. For myelin removal, the pelleted homogenate was then resuspended in 6 mL of 37% isotonic Percoll, and then underlayed with 5 mL of 70% isotonic Percoll in a 15 mL centrifuge tube. Tubes were then centrifuged at 600 x g for 40 min at 16-18°C, with no acceleration or deceleration. The top layer of myelin was removed using a 10 mL pipette and cells at the 37%–70% Percoll interphase were then recovered to 15 mL tubes and washed once in staining buffer. Cells were then incubated in FACs buffer on ice with CD16/CD32 blocking antibody for 15 min and incubated for 30 min with the corresponding primary antibody. Fluorescence Minus One (FMO), single-color and only cells controls were used in all the experiments for fluorescence compensation and gating strategy. After washing, cells were resuspended in staining buffer and acquired with LSRFortessa or FACSCanto II cytometers (BD Biosciences). To collect samples for subsequent bulk RNA-seq or sc-RNA-seq, samples were purified by Fluorescence-activated cell sorting (FACS) using a BD FACSAria IIu cell sorter (BD Biosciences). Pulse processing and cell viability dyes were used to exclude cell aggregates and dead cells. Data was analyzed using FlowJo v10.0 (Treestar, OR).

**List of flow cytometry antibodies**
| Antibody | IF dilution | Provider | Reference |
| --- | --- | --- | --- |
| CD74-AF647 | 1:1000 | BioLegend | 151003 |
| CX3CR1-APC/Fire750 | 1:300 | BioLegend | 149039 |
| CD45-PE/Cy7 | 1:800 | Invitrogen | 25-0451-81 |
| CD11b-PerCP-Cy5.5 | 1:200 | eBioscience | 45-0112-82 |

For bulk RNA-seq analysis, cells were recovered in Trizol LS reagent (10296010, Thermo Fisher), and RNA was extracted using the PicoPure RNA isolation Kit (ThermoFisher) according to manufacturer’s instruction. For single-cell RNA-seq analysis cells were recovered 1X PBS containing 0.04% BSA, and diluted to a final concentration of 7 x 10^5^ cells/ml.

### Multiplex immunoassay for detection of cytokines

The FirePlex-96 key cytokines (mouse) immunoassay panel (ab235656, Abcam) was used for unbiased detection of 17 different murine cytokines in brain tissue lysates from tumor-naive immunocompromised mice and H2030-BrM intracardially injected mice vehicle– or ibudilast-treated. For generation of brain tissue lysates from tumor-naive and metastatic mouse brains, brain metastasis were microdissected according to BLI signal and immediately snap-frozen in liquid nitrogen. Frozen tissue was subsequently homogenized and reconstituted in 1× Cell Lysis Buffer (9803, Cell Signaling Technology) plus protease inhibitors. Lysates were centrifuged for 20 min at 14,000 × g, and supernatants were used for the multiplex immunoassay at a concentration of 250 μg/mL.

### Detection of IFN-γ by ELISA

Lysis buffer (Cell Signaling ref. 9803S) with the following protease inhibitors: 200 mM Na3VO4, 500 mM NaF, 100 mM PMSF, was used to extract total protein. Protein lysates were obtained from wild-type C57BL/6 or intracranially injected mice with B16/F10-BrM cells and brain slices from wild-type athymic nude mice or intracardially injected with H2030-BrM cells. Tumors were obtained by dissecting Luciferase-tissue immediately adjacent to Luciferase+ cancer cells. Tissue was mechanically desegregated with the FastPrep-24™ 5G lysis system (MPBiomedical) by using zirconium beads at 6.0 m/s for 15 s followed by 10 min incubation on ice before lysis. For protein quantification, BCA protein color kit was used (Fisher Scientific, ref. 23227). The Invitrogen novex IFN-γ mouse ELISA Kit (Life technologies, KMC4021) was used according to the manufacturer’s instruction.

### RNA isolation and qRT-PCR

Whole RNA was isolated using the RNAeasy Mini Kit (QIAGEN). RNA from brain-metastatic cell lines (H2030-BrM and B16/F10-BrM cells containing shCTRL or sh*MIF*/sh*Mif*) was obtained from a confluent well from a six-well plate. RNA (1,000 ng) was used to generate complementary DNA (cDNA) using the iScript cDNA Synthesis Kit (1708890, Bio-Rad). Gene expression was analyzed using SYBR green gene expression assays (GoTaq qPCR Master Mix, A6002, Promega). The relative gene expression was normalized to a ‘housekeeping’ gene. The following primers were used (5’→3’, forward; reverse):

*B2M* (AGATGAGTATGCCTGCCGTG; TCATCCAATCCAAATGCGGC)

*B2m* (GACCGGCCTGTATGCTATCC; CAGTAGACGGTCTTGGGCTC)

*ACT* (CAAGGCCAACCGCGAGAAGAT; CCAGAGGCGTACAGGGATAGCAC)

*Act* (GGCACCACACCTTCTACAATG; GTGGTGGTGAAGCTGTAGCC)

*MIF* (TCTGCCATCATGCCGATGTT; TTGCTGTAGGAGCGGTTCTG)

*Mif* (CTTTGTACCGTCCTCCGGTC; CGTTCGTGCCGCTAAAAGTC)

Quantitative PCR reaction was performed on QuantStudio 6 Flex Real-Time PCR System (Applied Biosystems) and analyzed using the software QuantStudio 6 and 7 Flex software.

### Bulk RNA sequencing

Total RNA samples were processed with the “NEBNext Single Cell/Low Input RNA Library Prep” kit (NEB #E6420) by following manufacturer instructions. Briefly, an oligo(dT) primed reverse transcription with a template switching reaction was followed by double stranded cDNA production by limited-cycle PCR. (Non-directional) sequencing libraries were completed with the “NEBNext Ultra II FS DNA Library Prep Kit for Illumina” (NEB #E7805) and subsequently analysed on Illumina NextSeq 550 (using v2.5 reagent kits) in single-read fashion (85 bases) by following manufacturer’s protocols. Image analysis, per-cycle basecalling and quality score assignment was performed with Illumina Real Time Analysis software. Conversion of BCL files to FASTQ format was performed with Local Run Manager Generate FASTQ Analysis Module (Illumina).

Mouse reads were analysed with the Nextpresso ^84^ pipeline as follows: sequencing quality was checked with FastQC v0.11.0. Reads were aligned to the mouse genome (GRCm39) with TopHat-2.0.10^85^ using Bowtie 1.0.0^86^ and Samtools 0.1.19 ^87^, allowing 3 mismatches and 20 multihits. The Gencode vM26 gene annotation for GRCm39 was used. Read counts were obtained with HTSeq ^88^. Differential expression and normalization were performed with DESeq2 ^89^, filtering out those genes where the normalized count value was lower than 2 in more than 50 % of the samples. From the remaining genes, those that had an adjusted p-value below 0.05 FDR were selected. GSEAPreranked ^90^ was used to perform gene set enrichment analysis for several gene signatures on a pre-ranked gene list, setting 1000 gene set permutations. Only those gene sets with significant enrichment levels (FDR q-value < 0.25) were considered.

Pipeline used to preprocess RNA-seq data is available at GitHub: https://github.com/osvaldogc/nextpresso1.9.2. This section does not report original code.

## Survival signature

From the deregulated genes identified by the bulk RNA-seq, 151 had orthologues in the human genome, corresponding to 153 unique Ensembl values. 128 of these IDs had mRNA expression estimates in a 45 breast-cancer brain metastasis (BCBM) patients cohort previously described ^54^, and were further used to build a predictive gene signature model. “glmnet” R package was used to construct a prognostic model by least absolute shrinkage and selection operator (LASSO) Cox regression. The penalty regularization parameter lambda was selected through 10-times cross validation using the Harell C index (function cv.glmnet, family = “cox”, type.measure = “C”, set.seed(123) for reproducibility). Using the minimum deviance parameter lambda.min, 37 genes with coefficients higher or lower than 0 were selected for GS1, and 23 genes were selected for GS2. A risk score model was constructed based on the mRNA expression of each selected gene (log2CPMTMM normalized values) together with the coefficient generated by LASSO Cox regression ^91^.

The risk score was calculated for each patient, and the cohort was divided into high-risk and low-risk groups based on the median value of the risk score. Kaplan-Meier (KM) survival curves were plotted to visualize the association between the risk model and survival post brain metastasis (SPBM).

The association between the risk score model constructed using the cohort of 45 patients and SPBM was validated further by extending the initial cohort with 17 additional BCBM patients (newly sequenced, not previously published) (n = 62).

### Single cell RNA-seq library preparation and data analysis

FACS-purified CD45⁺ cells were processed for library preparation using the Chromium Next GEM Single Cell 3’ GEM, Library & Gel Bead Kit v3.1 (10x Genomics, PN-1000121), following the manufacturer’s instructions. The protocol included GEM generation, barcoding, GEM-RT clean-up, cDNA amplification, and library construction. The sequencing was conducted on an Illumina NextSeq 550 instrument equipped with v2.5 reagent kits, employing a PairEnd run type of 28 bp R1 and 56 bp R2. Primary data processing, including image analysis, per-cycle basecalling, and quality score assignment, was carried out using Real Time Analysis software (RTA v2, Illumina). Additionally, the conversion of BCL files to FASTQ format was executed with bcl2fastq2 within the Local Run Manager RNAfusion Analysis Module v2 (Illumina).

Seven 10x barcoded sequencing libraries were generated using the 10x Single Cell 3’ v3 kits (10x Genomics, Inc.). These libraries were sequenced in multiple rounds until each sample achieved a mean depth of 25,000 reads per cell. The mean number of reads per cell was calculated by processing the accumulated raw FASTQ files from each sequencing round with CellRanger Count (10X Genomics Cell Ranger v7.1.0) using default parameters. The sequencing data were aligned to the prebuilt 10x mouse GRCh38 (GENCODE v32/Ensembl 98) scRNA-seq reference genome (mm10-2020-A). This analysis yielded a median of 264 million sequenced reads per sample, with individual sample reads as follows: CTR_BrM: 264,577,836; BrM_VEH: 236,382,953; BrM_IBU: 295,303,588; CTR_MS: 96,922,258; MS: 222,018,909; CTR_AD: 335,324,101; AD: 275,467,604.

To remove potential contamination with human environmental RNA, the raw FASTQ data from the gene expression libraries of all samples were aligned to the human GRCh38 (GENCODE v32/Ensembl 98) and mouse GRCh38 (GENCODE v32/Ensembl 98) 10x prebuilt scRNA-seq reference genome (GRCh38_and_mm10-2020-A). This alignment was processed using CellRanger Count (10X Genomics Cell Ranger v7.1.0) with default parameters.

In total, 12,219 cells were retrieved from CTR_BrM, 9,445 from BrM_VEH, 11,850 from BrM_IBU, 29,818 from CTR_AD, 11,388 from AD, 13,189 from CTR_MS, and 18,849 from MS. Background ambient RNA and barcode-swapped reads were subsequently removed from the count matrix using the Cellbender v0.3.0 Python package ^92^ (Python v3.7.12) with default parameters. This processing yielded a refined count matrix, identifying barcodes corresponding to genuine cells.

After applying Cellbender, the cell counts were adjusted to 10,237 in CTR_BrM, 9,452 in BrM_VEH, 11,383 in BrM_IBU, 12,143 in CTR_AD, 10,262 in AD, 3,665 in CTR_MS, and 8,530 in MS. The removal of doublet artifacts was performed using the Scrublet v0.2.3 Python package ^93^ (Python v3.7.10) with default parameters. Following doublet removal, the final cell counts were 10,159 in CTR_BrM, 9,379 in BrM_VEH, 11,370 in BrM_IBU, 12,143 in CTR_AD, 9,283 in AD, 3,659 in CTR_MS, and 8,528 in MS.

All subsequent single-cell transcriptomic analyses were conducted using R Statistical Software (v4.2.0) with the Seurat v5 (v4.9.9.9059) package ^94^ (R: The R Project for Statistical Computing). For quality control, human genes and those expressed in fewer than three cells were excluded. After filtering out human genes, cells with more than 5% mitochondrial content or fewer than 1,500 detected mouse genes were also excluded. This filtering resulted in a total of 43,245 high-quality cells, distributed as follows: 7,360 in CTR_BrM, 5,202 in BrM_VEH, 5,870 in BrM_IBU, 9,108 in CTR_AD, 6,363 in AD, 2,885 in CTR_MS, and 6,457 in MS.

For each sample, UMI count data were normalized using a regularized negative binomial regression approach via SCTransform (sctransform v0.3.5). SCTransform was executed with the parameter vst.flavor set to “v2” to enhance regularization, and the method was set to “glmGamPoi” to fit a generalized linear Gamma-Poisson model, thereby improving the speed of the learning procedure and enhancing computational efficiency. In the subsequent step, mitochondrial content was regressed out in a second non-regularized linear regression.

To ensure that SCTransform residuals were computed for the top variable genes across all samples, the SelectIntegrationFeatures function was utilized to identify the top 3,000 highly variable genes (HVGs). These features were then employed in the PrepSCTIntegration function to recompute residuals for any missing values across the selected HVGs, utilizing stored model parameters. Upon obtaining the residuals for the common HVGs, Principal Component Analysis (PCA) was conducted on the SCT-normalized data of each sample, focusing on the top 3,000 HVGs.

Integration anchors were identified using the FindIntegrationAnchors function, specifying SCT as the normalization method, the top 3,000 HVGs as anchor features, and employing reciprocal PCA (RPCA) for reduction over the top 30 PCA components, with k set to 20 for anchor selection.

The seven datasets were integrated into a single dataset using the IntegrateData function, applying the previously identified anchors, SCT normalization, and the top 30 dimensions for the anchor weighting procedure. To facilitate dimensionality reduction, a new PCA was performed on the integrated dataset, followed by a nonlinear dimensionality reduction using Uniform Manifold Approximation and Projection (UMAP) applied to the 30 most significant PCA dimensions.

A Shared Nearest-Neighbor (SNN) graph was constructed utilizing the FindNeighbors function, based on the 30 most significant principal components (PCs). Cell clusters were subsequently identified through the implementation of the FindClusters function with default parameters.

For differential expression analysis within the integrated object comprising multiple SCT models, the dataset was prepared using the PrepSCTFindMarkers function. Differentially expressed genes between identified clusters were determined via a Wilcoxon rank-sum test, implemented through the Seurat package’s FindAllMarkers function, with default parameters. This analytical approach culminated in the identification of 16 distinct immune subpopulations. Gene expression plots were generated using the FeaturePlot function.

Enrichment scores for the identified gene signatures were computed using the AddModuleScore function with default settings, facilitating a systematic assessment of gene expression patterns across clusters and conditions. The visualizations were enhanced with the “RdBu” gradient color scheme from the RColorBrewer package. To enhance the interpretability of the expression data, the maximum and minimum cutoff values for visualization were adjusted, emphasizing biologically relevant expression ranges.

### Sampling of Human Tissues

Human brain metastasis tissue and CSF were collected by CNIO Biobank as the backbone of a collaborative nation-wide multicenter cohort, RENACER, integrated by 19 different hospitals and coordinated from the CNIO Biobank. Written informed consent for each donor was collected from each patient included in this study and surplus diagnostic samples were shipped to CNIO in less than 24 hours from surgery, under controlled temperature and other preanalytical variables, to warranty homogeneity and quality of the cohort. All the studies were conducted in accordance with recognized ethical guidelines (Declaration of Helsinki) and were approved by our institutional review board (IRB; CEI PI 25_2020-3). Comprehensive clinical information was also collected by CNIO Biobank associated to the samples.

Post-mortem cortical brain tissue blocks from human MS patients and controls, as well as the associated clinical and neuropathological data, were supplied by the Multiple Sclerosis & Parkinsońs Tissue Bank at Imperial College (London, UK), funded by the Multiple Sclerosis Society of Great Britain and Northern Ireland, registered charity 207495 (Dr. F. de Castrós request approved by communication dated on 19 January, 2023). Alternate coronal slices or brain hemispheres were fixed in 4% paraformaldehyde (PFA) for ∼2 weeks and were then cryoprotected in 30% sucrose for ∼1 week and frozen in isopentane precooled on a bed of dry ice. Microtome sections (50 μm) were obtained from all the cortical blocks, containing gray and white matters for immunohistochemical analysis as previously described^95^. Blocks from MS patients with white matter lesions and controls were studied here, from individuals with no history of neuropsychiatric disease in either case. The use of the UK Multiple Sclerosis Tissue Bank by Dr. F. de Castrós group at Instituto Cajal-CSIC has been approved (code 073/2021) by the Ethics Committee of Consejo Superior de Investigaciones Científicas (CSIC, Madrid, Spain). Control samples were chosen among those available to best match age, sex and postmortem interval with MS cases.

### Patient-derived organotypic cultures (PDOCs)

A total of 26 brain metastases from patients with lung cancer (11 cases), breast cancer (7 cases), colorectal cancer (3 cases), uterine cancer (2 cases), melanoma (2 cases) and prostate cancer (1 case) were obtained from RENACER collaborating hospitals: Hospital 12 de Octubre (Madrid), Hospital La Princesa (Madrid), Complejo Hospitalario Universitario de Albacete, Hospital Universitario de Burgos, Hospital Rio-Hortega (Valladolid) and Hospital Universitario Bellvitge (Barcelona). PDOCs were generated as described previously^70^. Briefly, after neurosurgical resection, brain metastasis samples were directly collected in neurobasal media A supplemented with 1x B27 (17504-044, Gibco), 1x N-2 (17502-048, Gibco), 25 ng/mL EGF (E9644, Sigma-Aldrich), 25 ng/mL bFGF (13256029, Gibco), 10 ng/mL NRG1-b1/HRG-b1 (396-HB, R&D Systems), 100 ng/mL IGF1 (291-G1, R&D Systems), 100 IU/mL penicillin/streptomycin and 1 μg/mL amphotericin B. Tissue was embedded in 4% low-melting agarose and cut in 250 μm slices using a vibratome.

PDOCs were treated with either DMSO or 50 μM ibudilast for 3 days. The human brain slices were fixed with 4% paraformaldehyde (PFA) overnight at 4°C and a free-floating immunofluorescence was performed afterwards. Proliferation was evaluated by manually counting Ki67+ cells in 3 brain slices per condition, and 3 FOV captured per slice.

### Bulk RNA-sequencing data processing

Raw RNA-sequencing (RNA-seq) reads were processed as follows: we used FASTQC v0.12.1 to check the quality of the sequencing reads and BBDuk (BBMap v38.93) to remove the adapter sequences and read ends with Phred quality scores lower than 10, discard reads shorter than 20 bp. Trimmed reads were aligned to the human genome (GRCh38) with STAR v0.74.0 ^96^ and Samtools v1.14 ^87^. Finally, mapped reads were counted and aggregated to a matrix of gene-level counts with HTSeq v0.13.5 ^88^. Genes with less than 10 counts across samples were filtered out. RNA-seq based counts were normalized using a variance stabilizing transformation with DESeq2 v1.34.0 ^89^ and then corrected for batch variation while preserving variation associated with the primary tumour with limma v3.50.1 ^97^. Differential expression was performed using DESeq2 ^89^. We only considered 25 patients from the RENACER cohort from six different primary sites (lung, n=11; breast, n=7; melanoma, n=3; colorectal, n=3; prostate, n=1; uterus, n=1), who had patient-derived organotypic cultures (PDOCs) treated with ibudilast and available RNA-seq data.

### Definition of a 11-gene signature associated with response to ibudilast

To select genes that were differentially expressed between responder (n=16) and non-responder group (n=7), we applied an adjusted p-value below 0.05 FDR and identified 51 genes. The responder group comprised 9 intermediate– and 9 super-responders. We quantified a 51-gene signature score using the weighted mean method from decoupleR ^98^ v2.8.0. We assigned a positive unit weight to genes associated with favorable response and a negative unit weight to genes associated with non-response. Considering the original 51-gene signature of ibudilast response, we used a drop out inference approach to optimize our signature genes while retaining predictive value. First, we computed the scores 1000 times, by dropping a variable percentage of the signature genes each time. This percentage ranges between 5% and 95%. Second, by checking all the occurrences where a gene was present, we calculated the average Mean Square Error (MSE) to compare the actual predictions with the predictions from the original signature (full score with all genes). Third, we built our ranking on the basis that a gene removal has a bigger effect on the predictions (higher MSE is associated with a gene being most useful/important). To assess the stability of this ranking, we applied bootstrap on the average MSE of each gene creating 100 samples of size equal to the minimum number of occurrences of all genes. The different gene rankings derived from the bootstrap samples showed a nice overlap of the gene ranks. Finally, we reduced the original 51-gene signature to a final signature comprising the 11 top-ranked genes (i.e. dropping out 78% of the original signature genes), as it showed a reasonable compromise between the loss in performance (MSE=1.83) and the ability to predict ibudilast response (area under the curve [AUC]=0.89). Patients were stratified into responders and non-responders based on the median value of the final 11-gene signature score.

### Estimation of differential pathway activity

We used the Univariate Linear Model (ULM) method from decoupleR ^98^ v2.8.0 to infer pathway activities, based on the t-values of DEGs between non-responders and responders and the PROGENy pathway weighted interaction network ^99^ (14 pathways).

### MIF quantification in clinical samples

Brain metastases were obtained from the CNIO Biobank that previously received them from Hospital Universitario 12 de Octubre and Hospital La Princesa. All samples followed protocols approved by our IRB (CEI PI 25_2020-3) and the Institutional Review Board of the Department of Neuroscience, University of Turin. Written informed consent was signed by each patient included in this study. IHC was performed at the CNIO Histopathology Core Facility using standardized automated protocols.

### MIF Detection in CSF

CSF samples from eight noncancer patients were obtained from the Biobank of Hospital Universitario Virgen de la Macarena, and from patients with lung cancer brain metastasis (six cases), breast cancer brain metastasis (two cases), melanoma brain metastasis (one case), and brain metastasis with other primary tumors (two cases) were obtained from the CNIO Biobank that previously received them from Hospital Universitario 12 De Octubre, Complejo Hospitalario Universitario de Vigo, Hospital Virgen De La Salud de Toledo, Hospital Universitario Vall d’Hebron and Complejo Hospitalario Universitario Santiago De Compostela. All samples were in compliance with protocols approved by their respective IRB (B.0001601, CEI PI 25_2020-v2, and CEI PI 25_2020-3). Written informed consent was signed by each patient included in this study. MIF levels in patients’ CSF were measured by ELISA following the manufacturer’s instructions (Merck, RAB0360-1KT).

### Meta-analysis of our gene signature across publicly accessible human scRNA-seq datasets

The analysis of scRNA-seq data from murine models of BrM, AD, and MS identified a specific gene signature for CD74+ myeloid cells, comprising a total of 672 genes. To evaluate the relevance of this signature within human patient datasets for MS and AD, publicly available single-cell expression datasets corresponding to the studies by Jäkel et al. (2019) ^75^ for MS and Gabitto et al. (2024) ^74^ for AD were accessed, downloaded and reprocessed as follows.

The MS scRNA-seq expression matrix and the corresponding annotation table were imported into R using the read.delim function. Row names in the annotation table were standardized by replacing colons and slashes with periods to ensure uniformity. Subsequently, a Seurat object was created to encapsulate both the expression data and its associated metadata. This Seurat object was then segmented into a list of individual datasets based on unique identifiers found in the “orig.ident” metadata column. Each dataset underwent normalization and variance stabilization via the SCTransform function, utilizing the “glmGamPoi” method to enhance computational efficiency.

Following normalization, SCTransform residuals were computed for the top variable genes across all datasets. The top 3,000 HVGs were selected as integration features using the SelectIntegrationFeatures function and were subsequently utilized in the PrepSCTIntegration function to recompute residuals for any missing values across these 3,000 HVGs, leveraging the stored model parameters. Once the residuals for the common HVGs were obtained, PCA was performed on the SCT-normalized data for each dataset, focusing on the top 3,000 HVGs.

Integration anchors were then identified using the FindIntegrationAnchors function, specifying SCT as the normalization method, the top 3,000 HVGs as anchor features, and employing reciprocal PCA (RPCA) as the reduction method over the top 30 PCA components, with 20 neighbors (k) selected for anchor identification. The individual datasets were subsequently integrated into a single dataset using the IntegrateData function, with the previously established anchors, SCT normalization, and the top 30 dimensions utilized in the anchor weighting procedure. To facilitate dimensionality reduction, a new PCA was executed on the integrated dataset, followed by a nonlinear dimensionality reduction using UMAP applied to the 30 most significant PCA dimensions.

For the analysis of two single-nucleus RNA sequencing datasets from AD, specifically derived from the dorsolateral prefrontal cortex (DLPFC) and the middle temporal gyrus (MTG), data were imported from RDS files into Seurat objects using the readRDS function, ensuring that all relevant information was retained. The disease status for each sample was categorized as either “normal” or “dementia.” Both AD datasets underwent normalization using the LogNormalize method, applying a scaling factor of 10,000 to standardize gene expression levels across the datasets.

Finally, the enrichment score for the Cd74+ cell-specific gene signature was computed across the three human datasets using the AddModuleScore function with default parameters. Feature plots were generated using the FeaturePlot function to visualize the spatial distribution of the calculated module scores across the UMAP embeddings for the respective datasets. The visualizations were enhanced with the “RdBu” gradient color scheme from the RColorBrewer package, and cutoff values were judiciously adjusted to emphasize the relevant expression ranges, thereby improving the interpretability of the results of the meta-analysis.

### Quantification and statistics

Data were analyzed using GraphPad Prism 8 software (GraphPad Software). For comparisons between two experimental groups in datasets that followed a normal distribution, an unpaired or paired (depending on the data), two-tailed Student t test was used. For comparisons that did not follow a normal distribution, a Mann-Whitney U test were performed. For multiple comparisons, ANOVA test was performed. For survival curves, P values were obtained with log-rank (Mantel–Cox) two-sided tests. χ2 test was performed for the comparison of group proportions.

## Data availability

Bulk RNAseq data from CD11b^+^; CD74^+^ versus CD11b^+^; CD74^−^ have been deposited to Gene Expression Omnibus (GEO) with the dataset identifier GSE 294461.

scRNA-seq data from CD45^+^ cells obtained brain metastasis, MS and AD experimental models have been deposited to Gene Expression Omnibus (GEO) with the dataset identifier GSE293921.

## Acknowledgements

We thank all members of the Brain Metastasis Group for critical discussion of the manuscript; the CNIO Core Facilities for their excellent assistance; and Estefanía Sánchez-Jiménez (F.dC. lab at Instituto Cajal-CSIC) and the staff at the Animal Facilities of the Instituto Cajal-CSIC for their extraordinary help with animals of the cuprizone demyelinating murine model of MS. Special thanks to patients and their families that donate valuable materials to RENACER, and all site personnel, investigators, funders and industry partners who supported the generation of the data within this study. Post-mortem cortical brain tissue blocks from human MS patients and controls, as well as the associated clinical and neuropathological data, were supplied by the Multiple Sclerosis & Parkinsońs Tissue Bank at Imperial College (London, UK), funded by the Multiple Sclerosis Society of Great Britain and Northern Ireland, registered charity 207495. We thank J. Massagué (MSKCC) for some of the BrM cell lines. This study was funded by La Marató (201906-30-31-32) (M.V.), MICIU (SAF2017-89643-R, SAF2014-57243-R, SAF2015-62547-ERC to M.V.; PID2022-143110OB-I00 and RED2024-153909-E to F.dC.), H2020-FETOPEN (828972) (M.V.), Bristol-Myers Squibb-MRA Young Investigator Award (498103) (M.V.), Cancer Research Institute (Clinic and Laboratory Integration Program CRI Award 2018 (54545) (M.V.), LAB AECC 2019 (LABAE19002VALI) (M.V.), AECC Coordinados (PRYCO234528VALI) (M.V.), ERC CoG (864759) (M.V.), ERANET-TRANSCAN-3 (TRANSCAN2021-203) (M.V.) with funds from Instituto de Salud Carlos III/ NextGenerationEU/ PRTR (AC20/00114) and FC AECC (TRNSC213878VALI), CaixaResearch Health (HR23-00051) (M.V.), MINECO-Ramón y Cajal (RYC-2013-13365) (M.V.), MICIU/AEI/10.13039/501100011033 and ERDF (RTI2018-102260-B-I00) (J.P.L-A), Generalitat Valenciana (PROMETEO/2020/007) (J.P.L-A), Ramón Areces Foundation (CIVP20S10662 to E.O-P/M-J.A.; CIVP19S8163 to M.V.; CIVP19A5917 to F.dC.), Spanish Ministry of Science and Innovation Competitiveness MCIN/AEI/ 10.13039/501100011033 and by “ERDF A way of making Europe” (RTI2018-099267-B-I00 and PID2022-136698OB-I00) (A.S.), Basque Government grants (IT1473-22) (A.S.), Alzheimer Association award (AARG-NTF-24-1304352) (A.S.), Breast Cancer Ireland (18239A01) (L.Y); Research Ireland (19/FFP/6443) (L.Y), (20/FFP-P/8597) (D.V), (23/SPP/11783) (LY, DV), Breast Cancer Now (2021JulyPCC1460) (L.Y, D.V), (2019AugSF1310 with the generous support of Walk the Walk) (DV), MCIN/AEI/10.13039/501100011033 and the European Union ‘‘NextGenerationEU’’/Plan de Recuperación Transformación y Resiliencia, PRTR (RTI2018-099357-B-I00, PID2021-1279880B, and TED2021-131611B-I00) (J.A.E.), CIBERFES (CB16/10/00282) (J.A.E.), and Foundation Leducq (17CVD04) (J.A.E.), La Caixa International PhD Program Fellowship-Marie Sklodowska-Curie (LCF/BQ/DI17/11620028) (P.G-G) and (LCF/BQ/DI19/11730044) (A.P.-A.), AECC postdoctoral fellowship (POSTD19016PRIE) (N.P.), MINECO-Severo Ochoa PhD Fellowship (BES-2017-081995) (L.A.-E.), FPI-Severo Ochoa Fellowship PRE2018-083478 awarded by MICIU/AEI /10.13039/501100011033 and by European Social Funds (FSE invierte en tu futuro) (Y.M.-M.). M.V. is an EMBO YIP member (4053). The CNIC is supported by the Instituto de Salud Carlos III (ISCIII), the Ministerio de Ciencia, Innovación y Universidades (MICIU) and the Pro CNIC Foundation, and is a Severo Ochoa Center of Excellence (grant CEX2020-001041-S funded by MICIU/AEI/10.13039/501100011033). The Instituto de Neurociencias is a ‘Centre of Excellence Severo Ochoa’ (CEX2021-001165-S funded by MCIU/AEI/10.13039/501100011033. CNIO is supported by the ISCIII, the Ministerio de Ciencia e Innovación, and is a Severo Ochoa Center of Excellence (SEV-2015-0510).

## Competing interests

Authors declare no conflict of interest

## Figures and figure legends

**Supplementary Figure 1.**
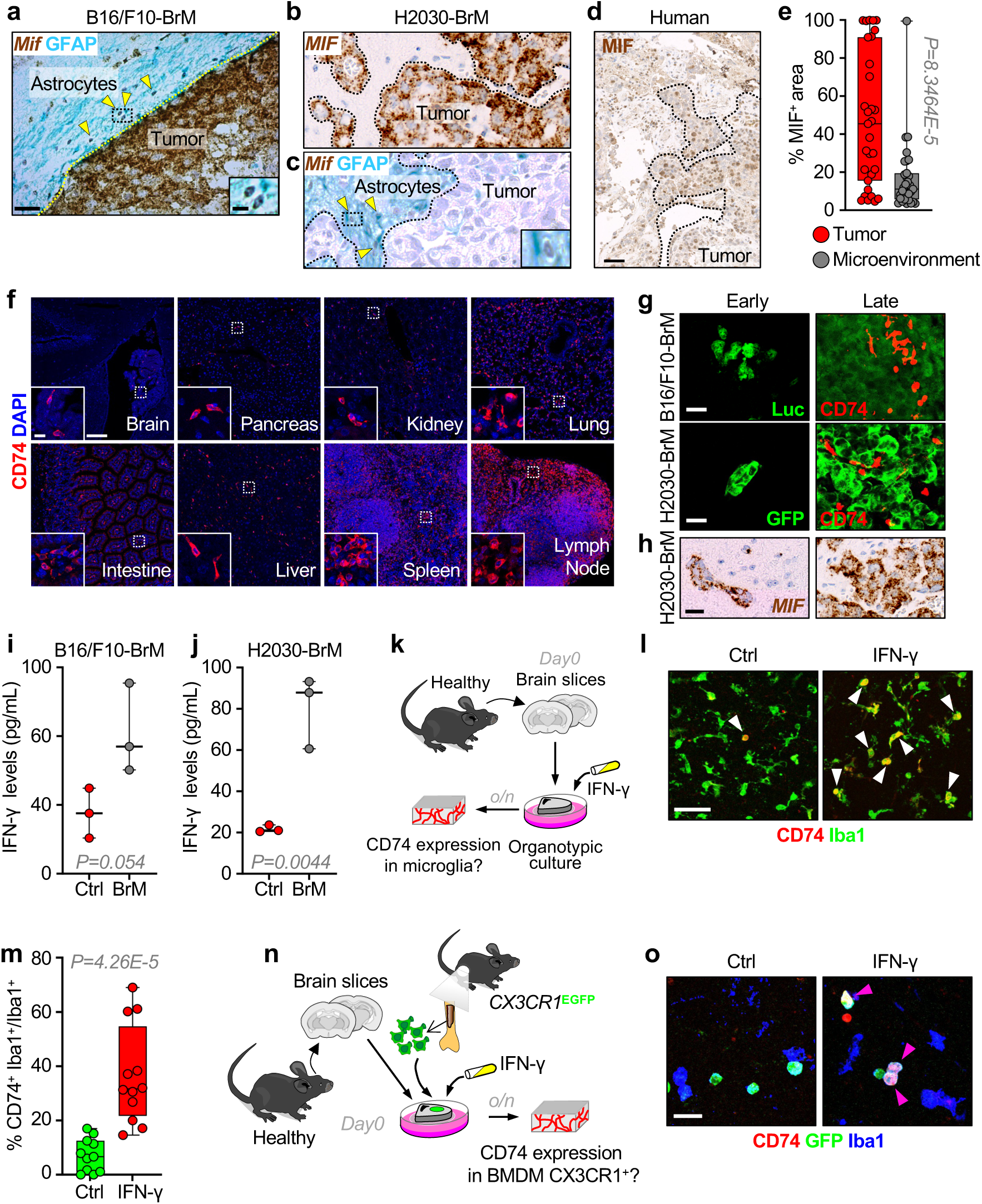
Cancer cells are the main source of MIF and CD74 expression in the brain tumor microenvironment is induced by IFN-γ. **a.** Representative immunohistochemistry of B16/F10-BrM brain metastasis with RNA *in situ* hybridization using RNAScope against *Mif* and GFAP immunohistochemistry. The yellow dotted lines delineate the tumoral area. The yellow arrows indicate astrocytes. The black dotted square highlight astrocytes with *Mif* expression. Scale bar: 20 µm. Magnification scale bar: 5 µm. **b.** Representative immunohistochemistry of H2030-BrM brain metastasis with RNA *in situ* hybridization using RNAScope against *MIF*. Black dotted lines delineate the tumoral area. Scale bar: 20 µm. **c.** Representative immunohistochemistry of H2030-BrM brain metastasis with RNA *in situ* hybridization using RNAScope against *Mif* and GFAP immunohistochemistry. The black dotted lines delineate the tumoral area. The yellow arrows indicate astrocytes. The black dotted square highlight astrocytes with *Mif* expression. Scale bar: 20 µm. Magnification scale bar: 5 µm. **d.** Representative immunohistochemistry staining against MIF in human brain metastasis. Scale bar: 50 µm. **e.** Quantification of immunohistochemical staining against MIF in multiple human brain metastases. The area MIF^+^ was calculated by normalizing the area expressing MIF in the tumoral or microenviromental compartment to the total area belonging to the tumor or microenvironment determined by H&E staining. Values are shown in a box-and-whisker plot where each dot represents one human sample, and the line corresponds to the median. n=31 human samples. *P* value was calculated using a paired two-tailed t-test. **f.** Immunofluorescence against CD74 in different organs from a healthy adult C56BL/6 mouse. White dotted squares outline the magnified area. Scale bar: 100 μm. Magnification scale bar: 10 μm. **g.** Immunofluorescences against CD74 and cancer cells (GFP in H2030-BrM, Luciferase [Luc] in B16/F10-BrM) at early (7 days in H2030-BrM and 3 days in B16/F10-BrM after intracardiac injection) and advanced stages (6 weeks in H2030-BrM and 14 days in B16/F10-BrM after intracardiac injection) of brain colonization. Scale bar: 25 μm. **h.** Representative immunohistochemistry with RNA *in situ* hybridization using RNAScope against *MIF* in tumors from H2030-BrM model at early (7 days after intracardiac injection) and advanced stages (6 weeks after intracardiac injection). **i.** Measure of IFN-γ levels (pg/mL) determined by ELISA in adult C57BL/6 healthy brains and B16/F10-BrM metastatic lesions. Values are shown in a dot plot where each dot represents a brain and the line corresponds to the median. *P* value was calculated using an unpaired two-tailed t-test. **j.** Measure of IFN-γ levels (pg/mL) determined by ELISA in brain slices derived from healthy immunocompromised mice and from brains with H2030-BrM established brain metastasis. Values are shown in dot plot where each dot represents a brain slice (H2030-BrM) and the line corresponds to the median. *P* value was calculated using an unpaired two-tailed t-test. **k.** Schema of the experimental design. Brain slices from a healthy adult C57BL/6 mouse were cultured overnight (o/n) with or without IFN-γ. **l.** Immunofluorescences against CD74 and Iba1 in adult healthy brain organotypic cultures stimulated or not with IFN-γ. Yellow arrows indicate the double-positive CD74^+^ Iba1^+^ cells. Scale bar: 50 μm. **m.** Quantification of l representing the percentage of CD74^+^ microglia. Each dot represents the mean percentage of cells per brain slice (n=12, 3 FOV quantified per brain slice) coming from 3 different mice from 2 independent experiments. Values are shown in a box-and-whisker plot where the line corresponds to the median. *P* value was calculated using an unpaired two-tailed t-test. **n.** Schema of the experimental design. Brain slices from a healthy adult C57BL/6 mouse were cultured with bone marrow cells extracted from *Cx3cr1*-EGFP mouse plated on top. Brain slices were cultured overnight (o/n) with or without IFN-γ. **o.** Immunofluorescences against CD74, Iba1 and GFP (CX3CR1) in adult healthy brain organotypic cultures with bone marrow-derived cells from a *Cx3cr1*-EGFP plated on top, cultured in the absence or presence of IFN-γ. Pink arrows indicate the triple-positive CD74+ Iba1+ CX3CR1+ cells. Scale bar: 25 μm.

**Supplementary Figure 2.**
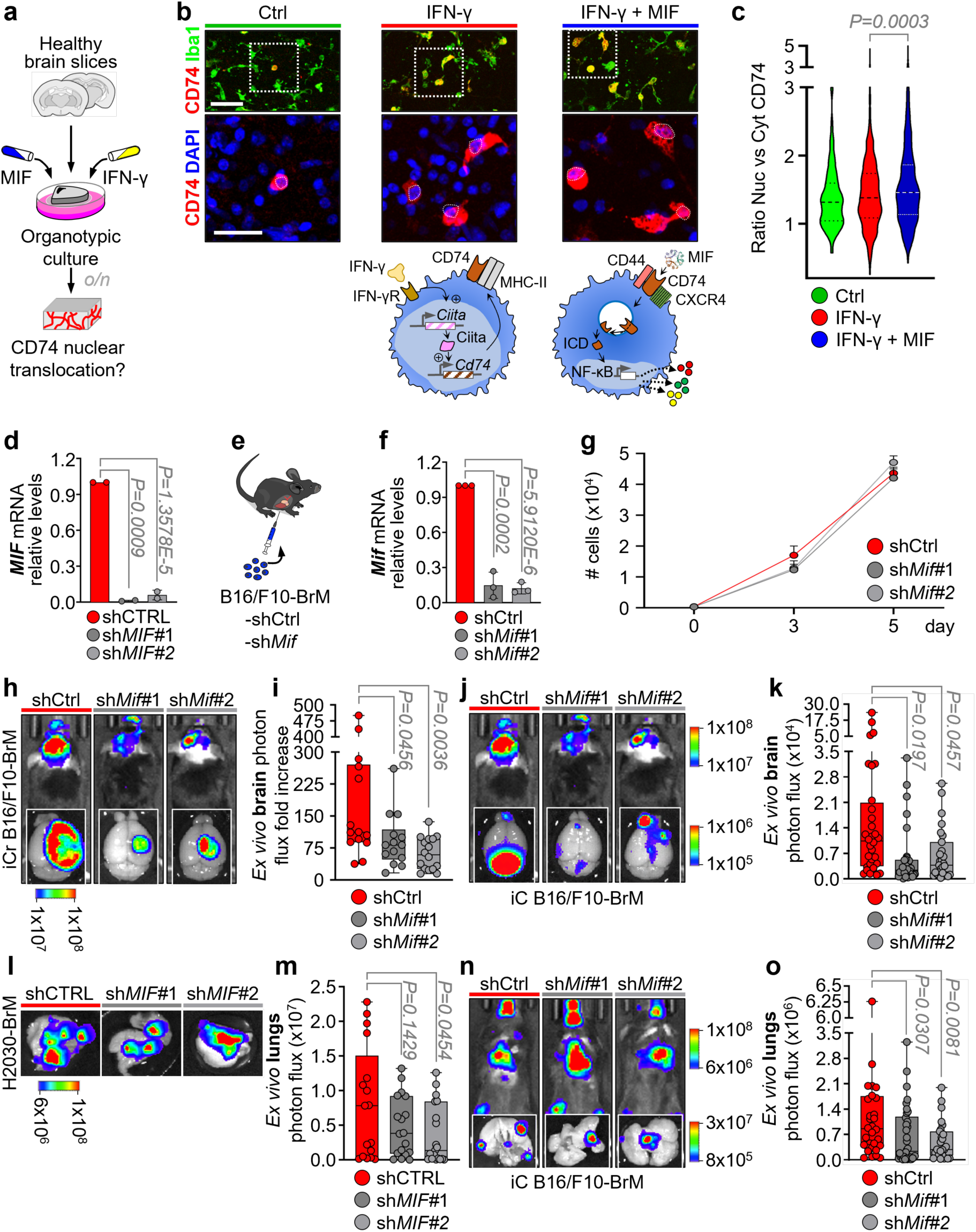
MIF induces CD74-ICD translocation to the nucleus and its silencing reduces B16/F10-BrM brain metastasis *in vivo*. **a.** Schema of the experimental design. Brain slices from a healthy adult C57BL/6 mouse were cultured overnight (o/n) with or without IFN-γ and MIF. **b.** Immunofluorescence staining for CD74, Iba1 and DAPI in brain organotypic cultures derived from healthy mice and subjected to the following stimulations: no treatment (left), IFN-γ (middle) or IFN-γ+MIF (right). White dotted squares indicate the area magnified in the panel below. Scale bar: 50 µm. The panel below show magnified images of the CD74+ Iba1+ cells. White dotted lines outline de DAPI^+^ nuclei. Scale bar: 25 µm. The schema below the immunofluorescence panel illustrates the hypothetical pathway CD74^+^ cells undergo under each treatment condition. Key functions of CD74 as a cytokine receptor have been shown to be dependent on CD44. CD74 additionally can complex with CXCR4 to facilitate MIF’s chemoattractant functions. **c.** Quantification of CD74-ICD found in the nucleus (DAPI) vs the membrane/cytoplasm on the magnified images shown in b. Values are shown in a Violin plot where each dot represents a cell and the line corresponds to the median. 100-600 cells were analyzed in 3 different brain slices per condition (4 FOV per brain slice) from 2 independent experiments. *P* value was calculated using an unpaired two-tailed t-test between the conditions IFN-γ vs IFN-γ+MIF. **d.** qRT-PCR of H2030-BrM cells for measurement of *MIF* expression after knockdown using shRNAs. Values are relative to the control and are shown in a dot plot where each dot represents a different technical replica from 2 different time points. **e.** Schema of the experimental design. B16/F10-BrM cells harboring a knockdown (KD) for MIF using two different shRNA were intracardially injected in wild-type mice. **f.** qRT-PCR of B16/F10-BrM cells for measurement of *Mif* expression after knockdown using shRNAs. Values are relative to the control and are shown in box-and-whisker plot where each dot represents a different technical replica from 3 different time points. **g.** Graph showing the growth of B16/F10-BrM cells *in vitro*. Proliferation was evaluated by plating equal numbers of cells at day 0, and taking pictures in the microscope at day 0, 3 and 5 and counting the cells at each time point. **h.** BLI *in vivo* images of mice injected with either B16/F10-BrM WT cells or KD for *MIF* 10 days after intracranial injection. *Ex vivo* BLI of brains (down) are also shown. **i.** Quantification of *ex vivo* BLI of brains from h normalized to *in vivo* BLI at day 3. Values are shown in a box-and-whisker plot where each dot represents a different animal and the line corresponds to the median. n=14 (shCtrl, *shMif#1*) and n=16 (*shMif#2*) mice from 2 independent experiments. *P* value was calculated using unpaired two-tailed t-test of each experimental condition against control. **j.** BLI *in vivo* images of mice injected with either B16/F10-BrM WT cells or KD for *MIF* 14 days after intracardiac injection. *Ex vivo* BLI of brains (down) are also shown. **k.** Quantification of *ex vivo* BLI of brains at day 14 (endpoint). Values are shown in a box-and-whisker plot where each dot represents a different animal and the line corresponds to the median. n=32 (shCtrl), n=34 (*shMif#1*), n=27 (*shMif#2*) mice from 3 independent experiments. *P* value was calculated using unpaired two-tailed t-test of each experimental condition against control. **l.** BLI *ex vivo* images of lungs from mice intracardially injected with either H2030-BrM WT cells or KD for *MIF* at endpoint (32-35 days after intracardiac injection). **m.** Quantification of *ex vivo* BLI of lungs from l. Values are shown in a box-and-whisker plot where each dot represents a different animal and the line corresponds to the median. n=17 (shCTRL), n=18 (*shMIF#1*), n=19 (*shMIF#2*) mice from 2 independent experiments. *P* value was calculated using an unpaired two-tailed t-test of each experimental condition against control. **n.** BLI *in vivo* and *ex vivo* images of lungs from mice intracardially injected with either B16/F10-BrM WT cells or KD for *Mif* at endpoint (14 days after intracardiac injection). **o.** Quantification of *ex vivo* BLI of lungs from n. Values are shown in a box-and-whisker plot where each dot represents a different animal and the line corresponds to the median. n=32 (shCtrl), n=34 (*shMif#1*), n=27 (*shMif#2*) mice from 3 independent experiments. *P* value was calculated using an unpaired two-tailed t-test of each experimental condition against control.

**Supplementary Figure 3.**
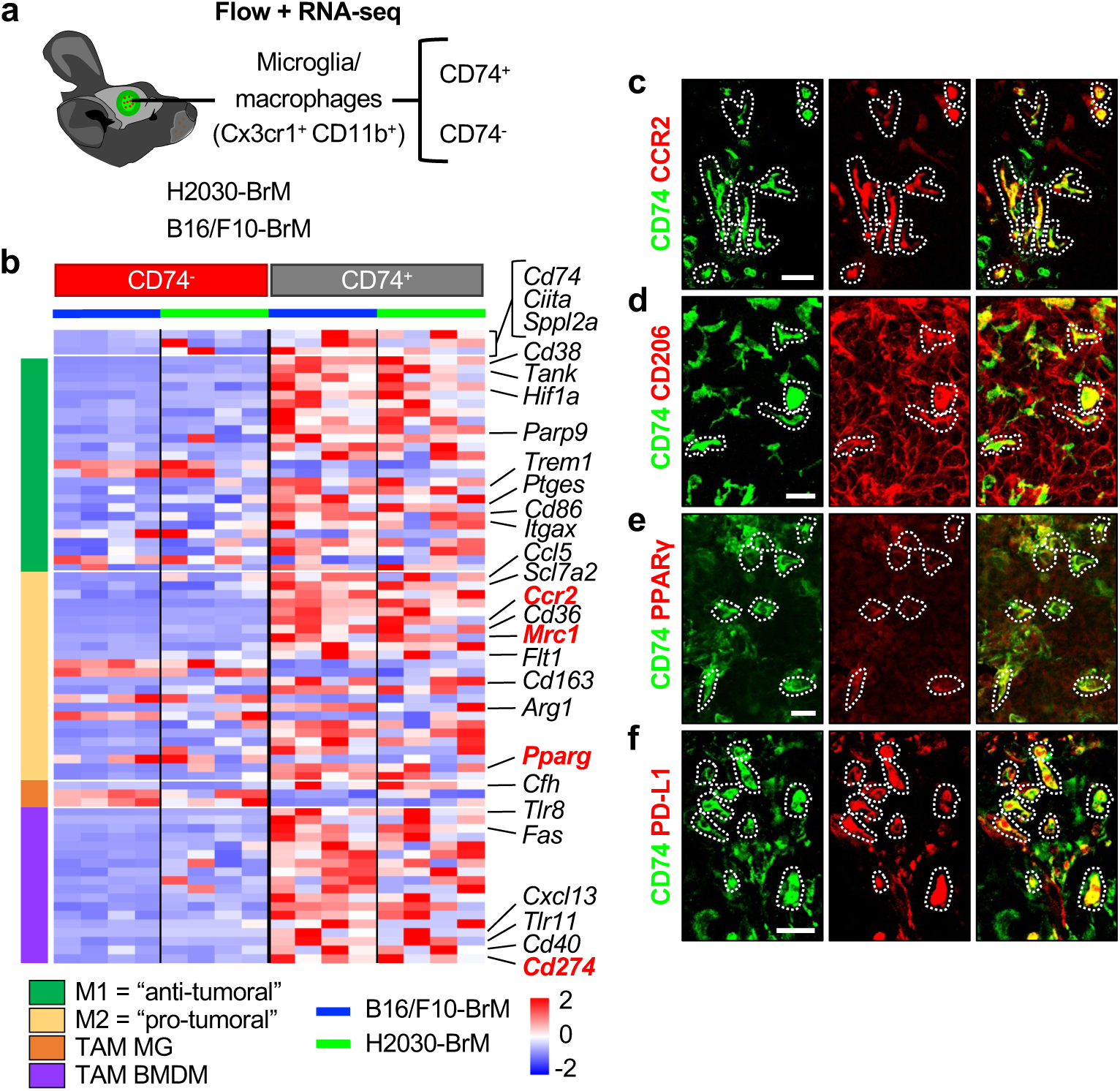
CD74^+^ microglia/macrophages are enriched in protumoral markers. **a.** Scheme of the experimental design. Microglia and macrophages (CX3CR1+ CD11b+) CD74+ or CD74-were isolated by fluorescence-activated cell sorting (FACS) from brain tumors derived from H2030-BrM or B16/F10-BrM inoculated mice and subjected to bulk RNA sequencing (RNA-seq) analysis. **b.** Heatmap of selected differentially expressed genes in CD74^+^ vs CD74^−^ microglia/macrophages. These genes include M1 and M2 markers, as well as tumor-associated microglia (TAM MG) and tumor-associated macrophages (TAM BMDM) markers that have been previously described in the literature. Genes written in red are the ones validated in the immunofluorescence panel at the right. *Sppl2a*, the gene encoding enzyme that catalyzes a crucial step in generating the CD74-ICD, as well as *Cd74* and its promoter regulator *Ciita* were enriched in the RNA-seq of CD74+ cells. **c. – f.** Immunofluorescence in metastatic lesions against CD74 and pro-tumoral markers shown in red in b. **c.** CCR2 (*Ccr2*), **d.** CD206 (*Mrc1*), **e.** PPAR γ (*Pparg*) **f.** PD-L1 (*Cd274*). Scale bar: 25 µm.

**Supplementary Figure 4.**
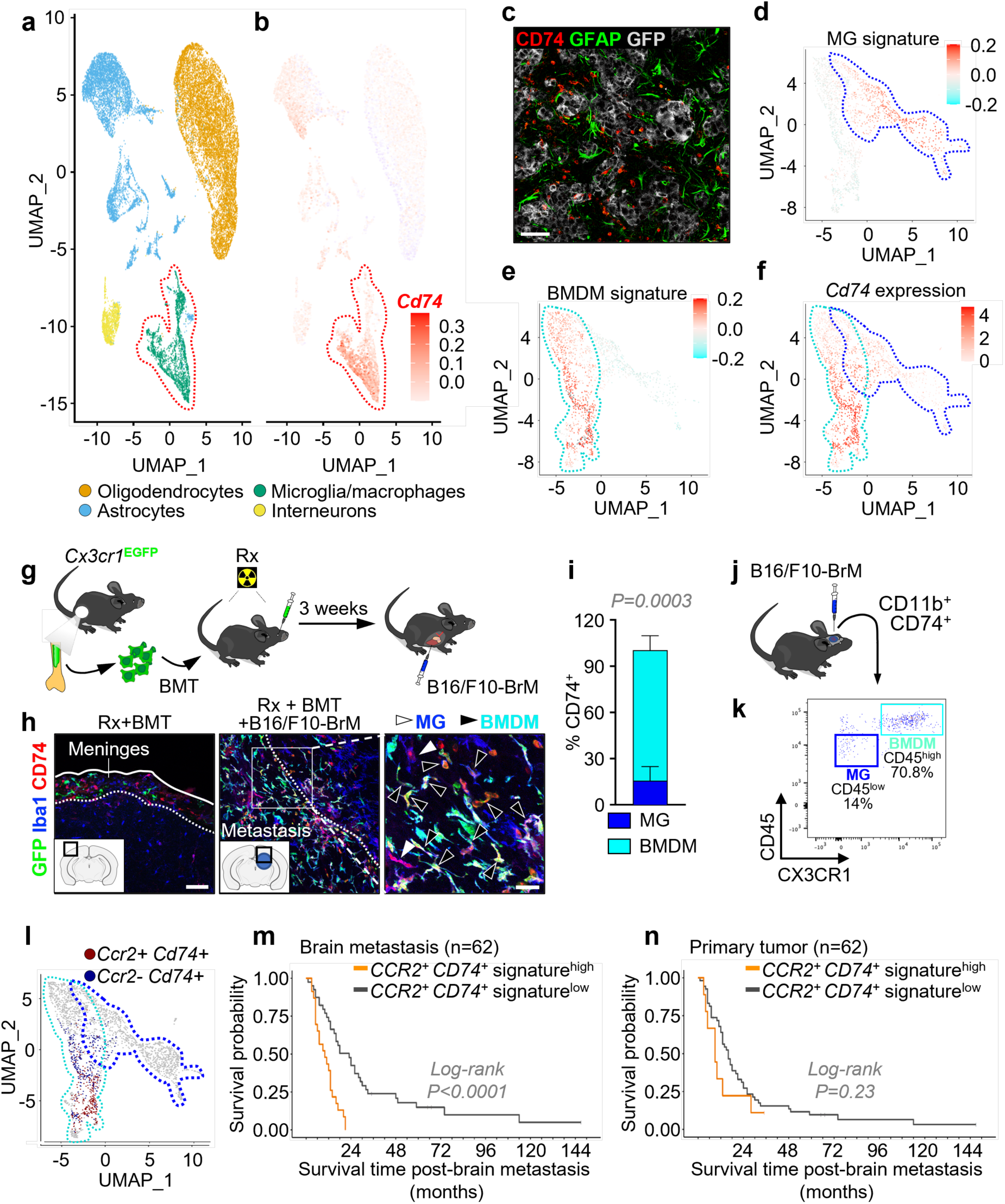
CD74^+^ are mainly bone marrow-derived macrophages with high expression of CCR2 and the *Cd74^+^ Ccr2^+^* signature has clinical relevance. **a.** Uniform manifold approximation and projection (UMAP) of populations found in the brain metastasis microenvironment of B16/F10-BrM model. **b.** UMAP showing *Cd74* expression, which is mostly expressed by the microglia/macrophages cluster shown in a. **c.** Immunofluorescence against GFP, CD74 and GFAP, labeling astrocytes, in H2030-BrM tumors. Scale bar: 50 µm. **d. –f.** Uniform manifold approximation and projection (UMAP) of the re-clusterized microglia/macrophage cluster in the single-cell RNA sequencing from a. **c.** Representation of the microglia (MG) signature. The area of expression is delineated by dark blue dotted lines. **d.** Representation of bone marrow-derived macrophage (BMDM) signature. The area of expression is delineated by light blue dotted lines. **e.** Representation of *Cd74* expression. **g.** Schema of the experimental design. Young adult C57BL/6 mice were irradiated with two doses 4.5 Gy. On the same day, bone marrow-derived cells were extracted from the femur and tibia of young adult C57BL/6 *Cx3cr1*-EGFP mice. The bone marrow-derived cells were retro-orbitally injected in the previously irradiated mice. 3 weeks after the bone marrow transplant (BMT), mice were intracardially injected with B16/F10-BrM cells. **h.** Immunofluorescence against CD74, Iba1 and GFP (CX3CR1) in brains from irradiated and bone marrow-transplanted mice. Left: no metastasis, white dotted line delineates the meninges. Middle: brain harboring B16/F10-BrM metastasis. White dotted line outlines the tumor border. White square outlines the area magnified in the right panel. Right: magnification showing CD74^+^ that are either Iba1^+^ CX3CR1^−^ microglia (MG, white arrows) and Iba1^+^ CX3CR1^+^ macrophages (BMDM, black arrows) in B16/F10-BrM tumor. Scale bar: 75 μm. Magnification scale bar 25 μm. **i.** Quantification of immunofluorescence in h. The bar represents the mean percentage out of the total CD74^+^ cells that are microglia (MG, dark blue) or macrophages (BMDM, light blue). n=6 brains from 2 independent experiments, with 8-14 FOV quantified per brain. *P* value was calculated using a paired two-tailed t-test. **j.** Schema of the experimental design. Mice were intracranially injected with B16/F10-BrM cells, and brains were subjected to flow cytometry analysis. **k.** Flow cytometry plot from B16/F10-BrM intracranially injected brains, the gating was done on CD11b^+^ CD74^+^ cells. Dark blue square delineates the microglia (MG, CD45^low^), while light blue square surrounds the macrophages (BMDM, CD45^high^). The graph represents a pool of 3 brains. **l.** Uniform manifold approximation and projection (UMAP) of the re-clusterized microglia/macrophage cluster in the single-cell RNA sequencing from a, representing the double positive *Ccr2*^+^ *Cd74*^+^ cells (red dots) and the *Ccr2*^−^ *Cd74*^+^ cells (blue dots). **m.** Survival graph of 62 patients with brain metastasis from breast cancer. Signature derived from *Cd74^+^ Ccr2^+^* cells was scored in RNA-seq data from brain metastasis human samples. The orange line represents patients enriched in the *Cd74^+^ Ccr2^+^* signature derived from the single-cell RNA sequencing, and the black line represents patients with lower enrichment of this signature. **n.** Survival graph of 62 patients with brain metastasis from breast cancer. Signature derived from *Cd74^+^ Ccr2^+^* cells was scored in RNA-seq data from breast primary tumors (matched samples from m). The orange line represents patients enriched in the *Cd74^+^ Ccr2^+^* signature derived from the single-cell RNA sequencing, and the black line represents patients with lower enrichment of this signature.

**Supplementary Figure 5.**
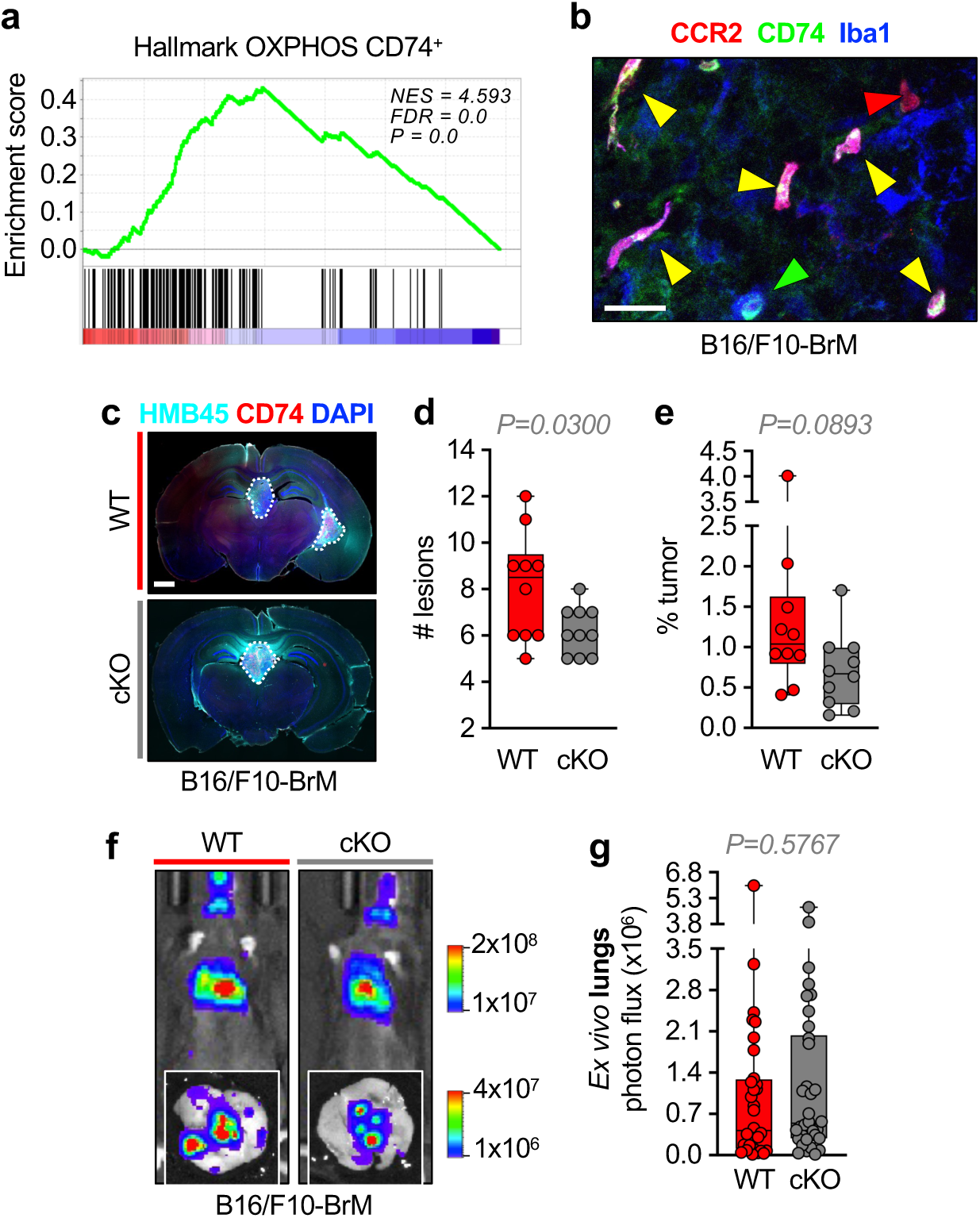
CD74^+^ cells are mainly BMDM with transcriptional enrichment in OXPHOS pathway. **a.** Gene set enrichment analysis of the oxidative phosphorylation (OXPHOS) that is enriched in the bulk RNA-seq of CD74^+^ microglia macrophages vs CD74^−^. **b.** Representative immunofluorescence in a B16/F10-BrM tumor against Iba1, CD74 and GFP (reporter of the Cre expression under the *Ccr2* promoter). The arrows label the microglia/macrophages (Iba1+) that are: CCR2^+^ CD74^+^ (yellow), CCR2^+^ CD74^−^ (red), CCR2^−^ CD74^+^ (green). Scale bar: 25 µm. **c.** Immunofluorescence against DAPI, HMB45 (melanoma marker) and CD74 in brain sections from WT and cKO^CCR2^-*Oma1* mice intracardially injected with B16/F10-BrM cells. The white dotted lines outline the tumoral lesions. Scale bar: 1 mm. **d.** Quantification of the total number of tumoral lesions per representative brain sections from WT and cKO^CCR2^-*Oma1* mice. Values are shown in a box-and-whisker plot where each dot represents a different animal and the line corresponds to the median. n=10 from 2 independent experiments. *P* value was calculated using an unpaired two-tailed t-test. **e.** Quantification of tumoral area per brain from representative brain sections. Percentage of tumoral area was calculated by measuring the total area belonging to the tumors normalized to the total area of the brain sections evaluated. Values are shown in a box-and-whisker plot where each dot represents a different animal and the line corresponds to the median. n=10 from 2 independent experiments. *P* value was calculated using an unpaired two-tailed t-test. **f.** Representative BLI *in vivo* images of WT and cKO CCR2-*Oma1* mice 14 days after intracardiac injection. *Ex vivo* BLI of lungs (down) are also shown. **g.** Quantification of *ex vivo* BLI of lungs from f. Values are shown in a box-and-whisker plots where each dot represents a different animal and the line corresponds to the median. n=33 (WT) and 35 (cKO) mice from 3 independent experiments. *P* value was calculated using an unpaired two-tailed t-test.

**Supplementary Figure 6.**
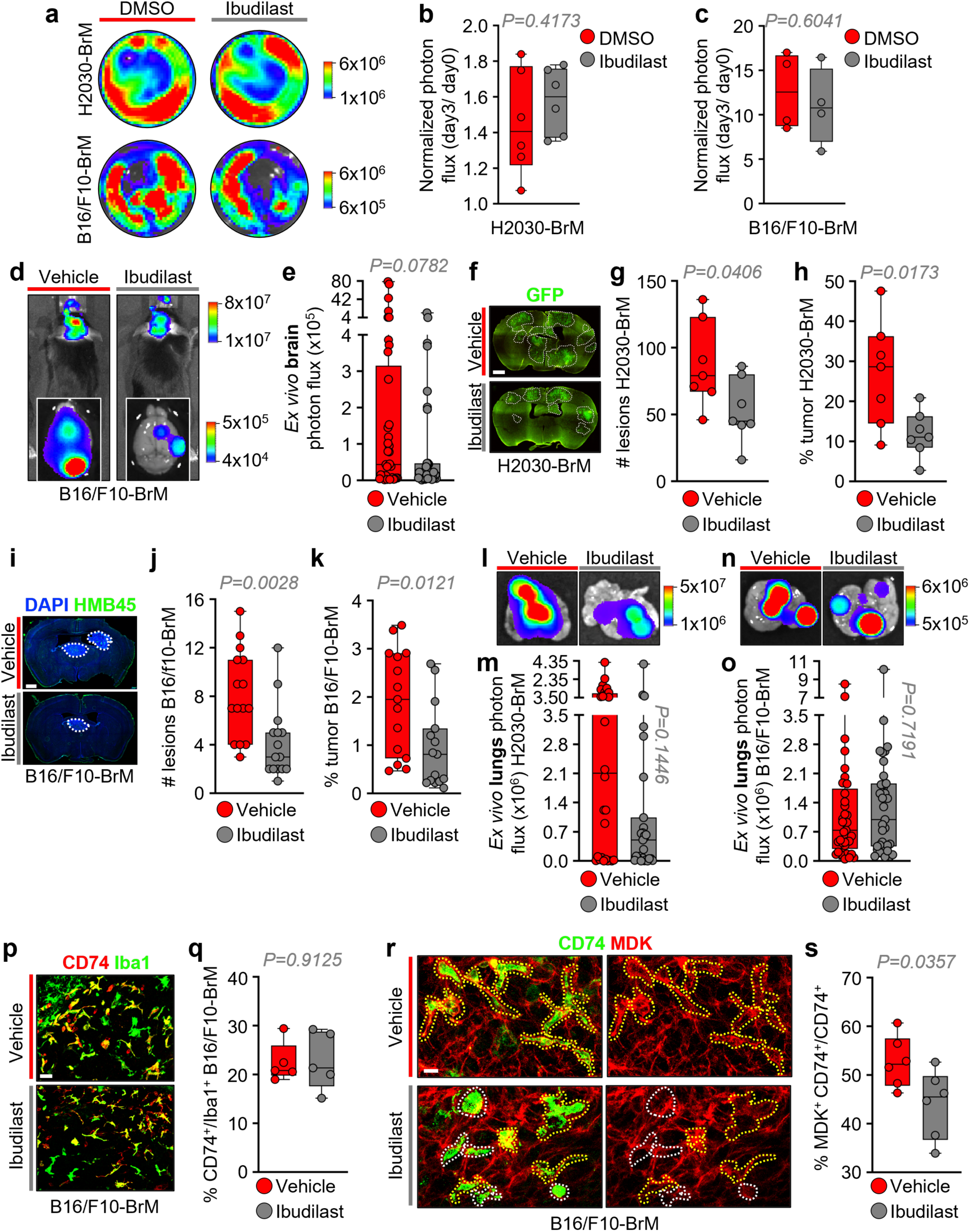
Ibudilast reduces brain metastasis *in vivo* without comprising the viability of cancer cells *in vitro*. **a.** Representative BLI images of *in vitro* cultured cancer cells H2030-BrM (top) and B16/F10-BrM (bottom) after 3 days of treatment with ibudilast (50 μM). **b. c.** Quantification of *in vitro* BLI of H2030-BrM cells (**b**) and B16/F10-BrM cells (**c**) from a measured at day 3 normalized to day 0. Values are shown in box– and-whisker plots where each dot represents a different well and the line corresponds to the median. n=4 wells in B16/F10-BrM cells and n=6 wells in H2030-BrM cells. *P* value was calculated using an unpaired two-tailed t-test. **d.** Representative BLI *in vivo* images of vehicle– or ibudilast-treated mice 14 days after intracardiac injection of B16/F10-BrM. *Ex vivo* BLI of brains are also shown. **e.** Quantification of *ex vivo* BLI of brains from d. Values are shown in a box-and-whisker plot where each dot represents a different animal and the line corresponds to the median. n=37 (vehicle) and 35 (ibudilast) mice from 4 independent experiments. *P* value was calculated using an unpaired two-tailed t-test. **f.** Representative immunofluorescence against GFP (labeling the H2030-BrM cells) in brain sections from vehicle– and ibudilast-treated mice intracardially injected with H2030-BrM cells. White dotted lines outline the tumoral lesions. Scale bar: 1 mm. **g.** Quantification of the total number of tumoral lesions per brain evaluated in representative brain sections. Values are shown in a box-and-whisker plot where each dot represents a different animal and the line corresponds to the median. n=7 mice per condition. *P* value was calculated using an unpaired two-tailed t-test. **h.** Quantification of tumoral area per brain from representative brain sections. Percentage of tumoral area was calculated by measuring the total area belonging to the tumors normalized to the total area of the brain sections evaluated. Values are shown in a box-and-whisker plot where each dot represents a different animal and the line corresponds to the median. n=7 mice per condition. *P* value was calculated using an unpaired two-tailed t-test. **i.** Representative immunofluorescence against DAPI and HMB45 (melanoma marker) in brain sections from vehicle– and ibudilast-treated mice intracardially injected with B16/F10-BrM cells. White dotted lines outline the tumoral lesions. Scale bar: 1 mm. **j.** Quantification of the total number of tumoral lesions per brain evaluated in representative brain sections. Values are shown in a box-and-whisker plot where each dot represents a different animal and the line corresponds to the median. n=15 mice per condition from 4 independent experiments. *P* value was calculated using an unpaired two-tailed t-test. **k.** Quantification of tumoral area per brain from representative brain sections. Percentage of tumoral area was calculated by measuring the total area belonging to the tumors normalized to the total area of the brain sections evaluated. Values are shown in a box-and-whisker plot where each dot represents a different animal and the line corresponds to the median. n=15 mice per condition from 4 independent experiments. *P* value was calculated using an unpaired two-tailed t-test. **l.** Representative BLI *ex vivo* images of lungs from vehicle– or ibudilast-treated mice at endpoint (32-35 days after intracardiac injection of H2030-BrM cells). **m.** Quantification of *ex vivo* BLI of lungs from l. Values are shown in a box-and-whisker plot where each dot represents a different animal and the line corresponds to the median. n=28 (vehicle) and 27 (ibudilast) mice from 3 independent experiments. *P* value was calculated using unpaired two-tailed t-test. **n.** Representative BLI *ex vivo* images of lungs from vehicle– or ibudilast-treated mice at endpoint (14 days after intracardiac injection of B16/F10-BrM cells). **o.** Quantification of *ex vivo* BLI of lungs from n. Values are shown in a box-and-whisker plot where each dot represents a different animal and the line corresponds to the median. n=37 (Vehicle) and 35 (Ibudilast) mice from 4 independent experiments. *P* value was calculated using an unpaired two-tailed t-test. **p.** Immunofluorescence against CD74 and Iba1 in B16/F10-BrM tumors from vehicle– and ibudilast-treated mice. Scale bar: 25 µm. **q.** Quantification of images shown in p. representing the percentage of CD74^+^ area normalized to the Iba1^+^ area. Values are shown in a box-and-whisker plot where each dot represents the mean percentage of CD74^+^ normalized to the Iba1^+^ area per brain, and the line corresponds to the median. n=5 mice, 4-8 FOV per mouse. *P* value was calculated using an unpaired two-tailed t-test. **r.** Immunofluorescence against CD74 and MDK in Iba1^+^ cells in B16/F10-BrM tumors from vehicle– and ibudilast-treated mice. White dotted lines outline CD74^+^ MDK^−^ cells, while yellow dotted lines outline double-positive CD74^+^ MDK^+^ cells. Images with the red-channel only (MDK) are shown for clarity. Evaluation of the expression was done in Iba1+ cells. Scale bar: 10 µm. **s.** Quantification of images shown in r, values are shown in a box-and-whisker plot where each dot represents the mean percentage of CD74^+^ Iba1^+^ cells positive for MDK normalized to the total CD74^+^ Iba1^+^ cells. 12 FOV were captured per brain, in n=6 brains from 3 independent experiments. *P* value was calculated using an unpaired two-tailed t-test.

**Supplementary Figure 7.**
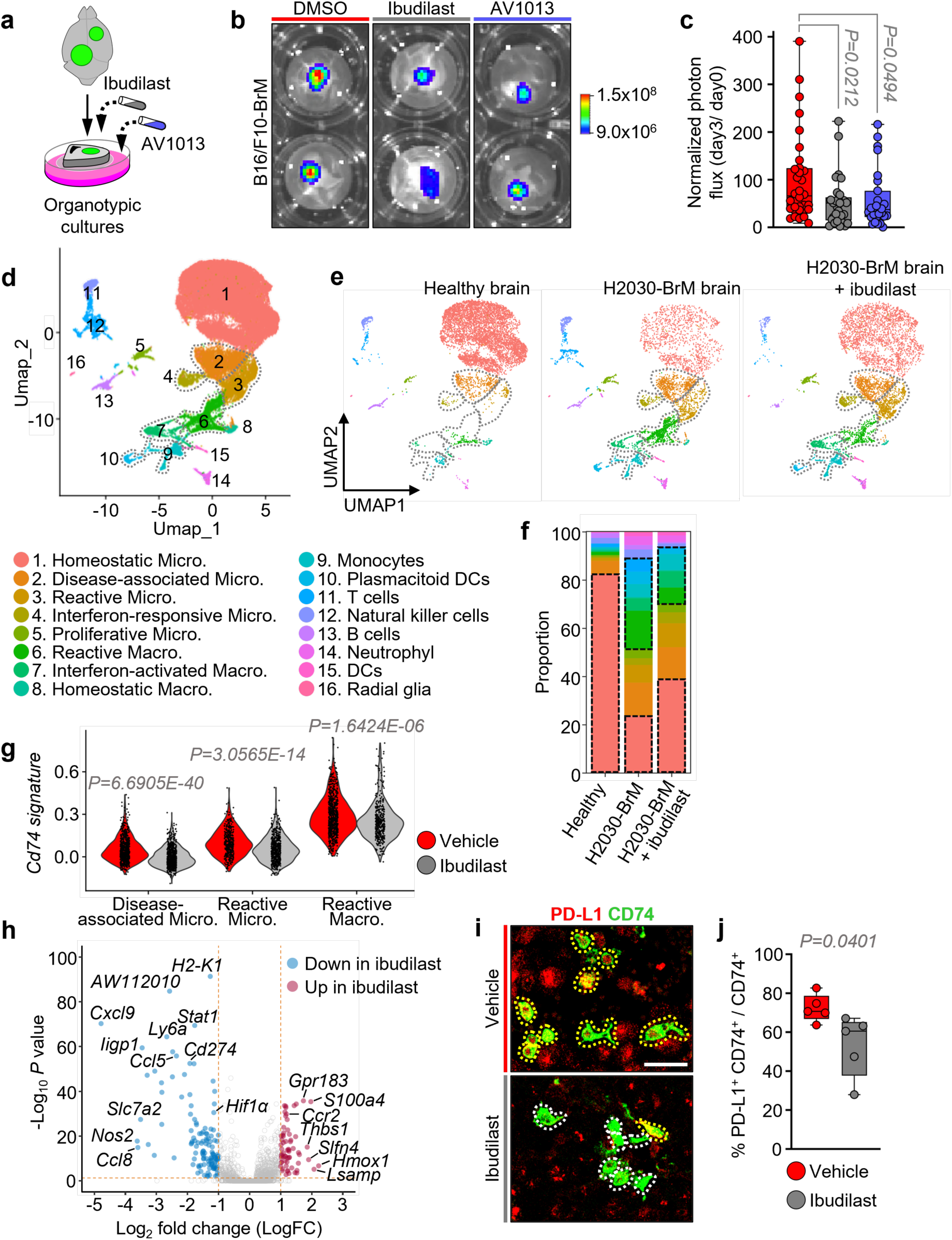
Ibudilast acts mainly through MIF inhibition and reverts the protumorigenic signature of CD74^+^ microglia/macrophages *in vivo*. **a**. Schema of the experimental design. Brain slices from B16/F10-BrM brain metastasis were cultured ex vivo (organotypic cultures) and ibudilast 50 μM or AV1013 50 μM was added and left for 3 days. BLI was measured at day 0 and day 3. **b**. Representative BLI images of brain slices with established brain metastasis from B16/F10-BrM at day 3 of the experiment. **c**. Quantification of *ex vivo* BLI of images in b, measured at day 3 normalized to BLI a day 0 of the experiment. Values are shown in a box-and-whisker plot where each dot represents a different slice, and the line corresponds to the median. n=32 (DMSO), n=29 (ibudilast) and n=27 (AV1013) from 2 independent experiments. *P* value was calculated using an unpaired two-tailed t-test of each experimental condition against control. **d**. Uniform manifold approximation and projection (UMAP) of CD45^+^ populations found in the healthy and brain metastasis microenvironment of H2030-BrM model, showing the existence of 16 clusters. Each cluster is represented with a unique color and number. Micro.: microglia, Macro.: macrophages, DC: dendritic cells. **e**. Uniform manifold approximation and projection (UMAP) of CD45^+^ populations by condition: left: healthy brain, middle: H2030-BrM metastatic brain and right: H2030-BrM metastatic brain treated with ibudilast. **f**. Stacked bar plot showing the proportional distribution of annotated immune and glial subpopulations based on single-cell RNA sequencing of CD45⁺ cells across experimental conditions: healthy, H2030-BrM untreated and ibudilast-treated brains. **g**. Violin plot presenting the distribution of *Cd74* signature score in the myeloid CD74^+^ populations (disease-associated microglia, reactive microglia and reactive macrophages) in vehicle– and ibudilast-treated mice with H2030-BrM metastasis. **h**. Volcano plot showing the deregulated genes in the reactive macrophages (cluster 6) from ibudilast-treated H2030-BrM mice to those treated with vehicle in H2030-BrM metastasis. **i**. Immunofluorescence against PD-L1 and CD74 in H2030-BrM tumors from vehicle– and ibudilast-treated mice. Scale bar: 25 µm. **j**. Quantification of images shown in i. Values are shown in a box-and-whisker where each dot represents the mean percentage of CD74^+^ Iba1^+^ cells positive for PD-L1 normalized to the total CD74^+^ Iba1^+^ cells per brain. 25-50 FOV were captured per brain, in n=5 brains. *P* value was calculated using an unpaired two-tailed t-test.

**Supplementary Figure 8.**
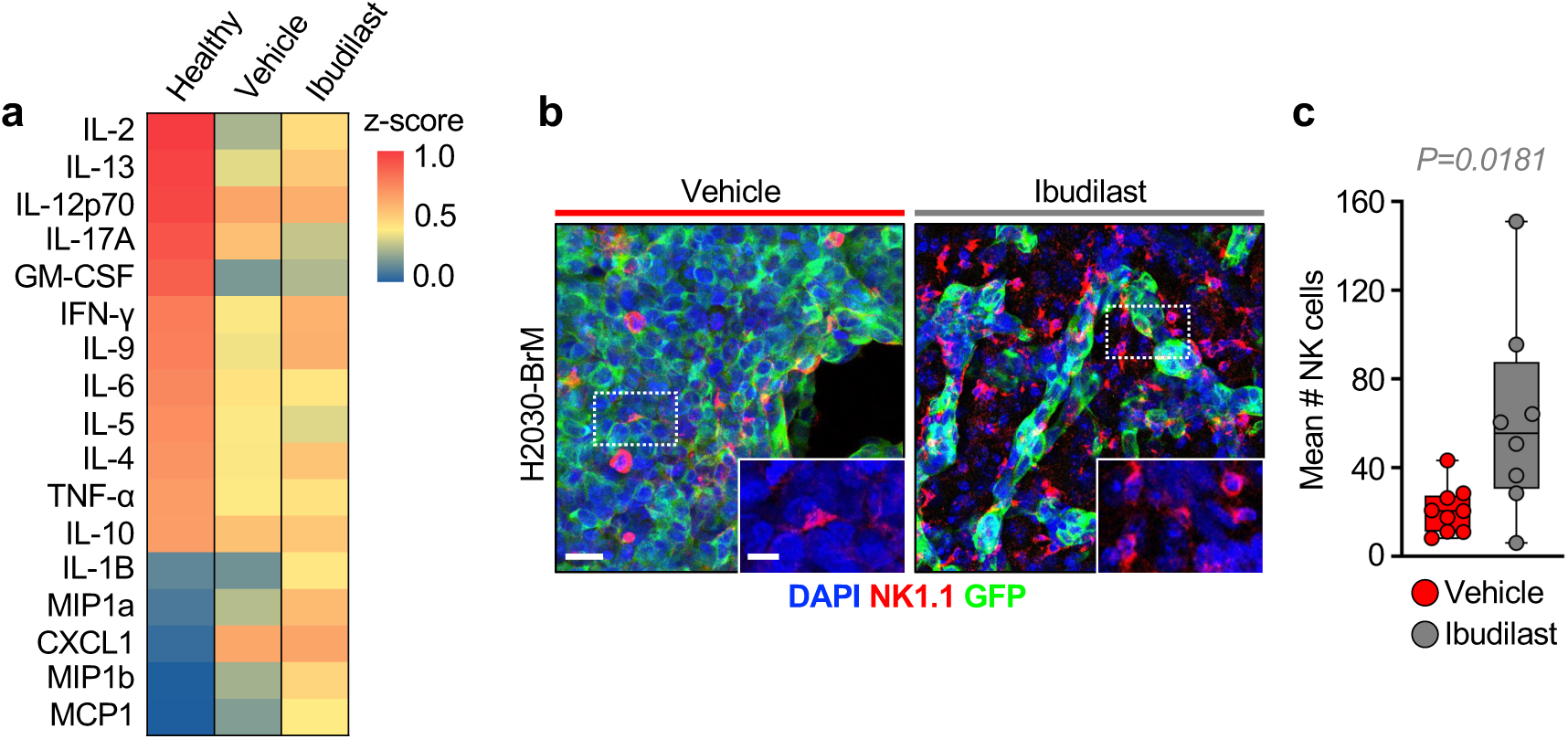
Ibudilast-treated brains show a recovery in IL-2 levels, which correlates with higher infiltration of natural killer cells. **a.** Quantification of 17 murine cytokines by multiplex cytokine assay in tumor-naïve (healthy) brains and brains with H2030-BrM metastasis treated with vehicle or ibudilast. Cytokine levels were measured by multiplex cytokine assay, and results are displayed as a heatmap of z-score–transformed mean concentrations (n=3, normal brains; n=5, brain metastasis brains). **b.** Immunofluorescence against DAPI, NK1.1 and GFP (labeling H2030-BrM cells) in tumors from vehicle– or ibudilast-treated mice. Scale bar: 25 μm. Magnification scale bar: 10 μm. **c.** Quantification of images shown in b, assessing the mean number of NK cells per FOV. Values are shown in a box-and-whisker plot where each dot represents the mean number of NK cells per FOV in brain lesions and the line in the box corresponds to the median. n=9 (vehicle) and n=8 (ibudilast) brains, with 4 FOV per brain from 2 independent experiments. *P* value was calculated using an unpaired two-tailed t-test.

**Supplementary Figure 9.**
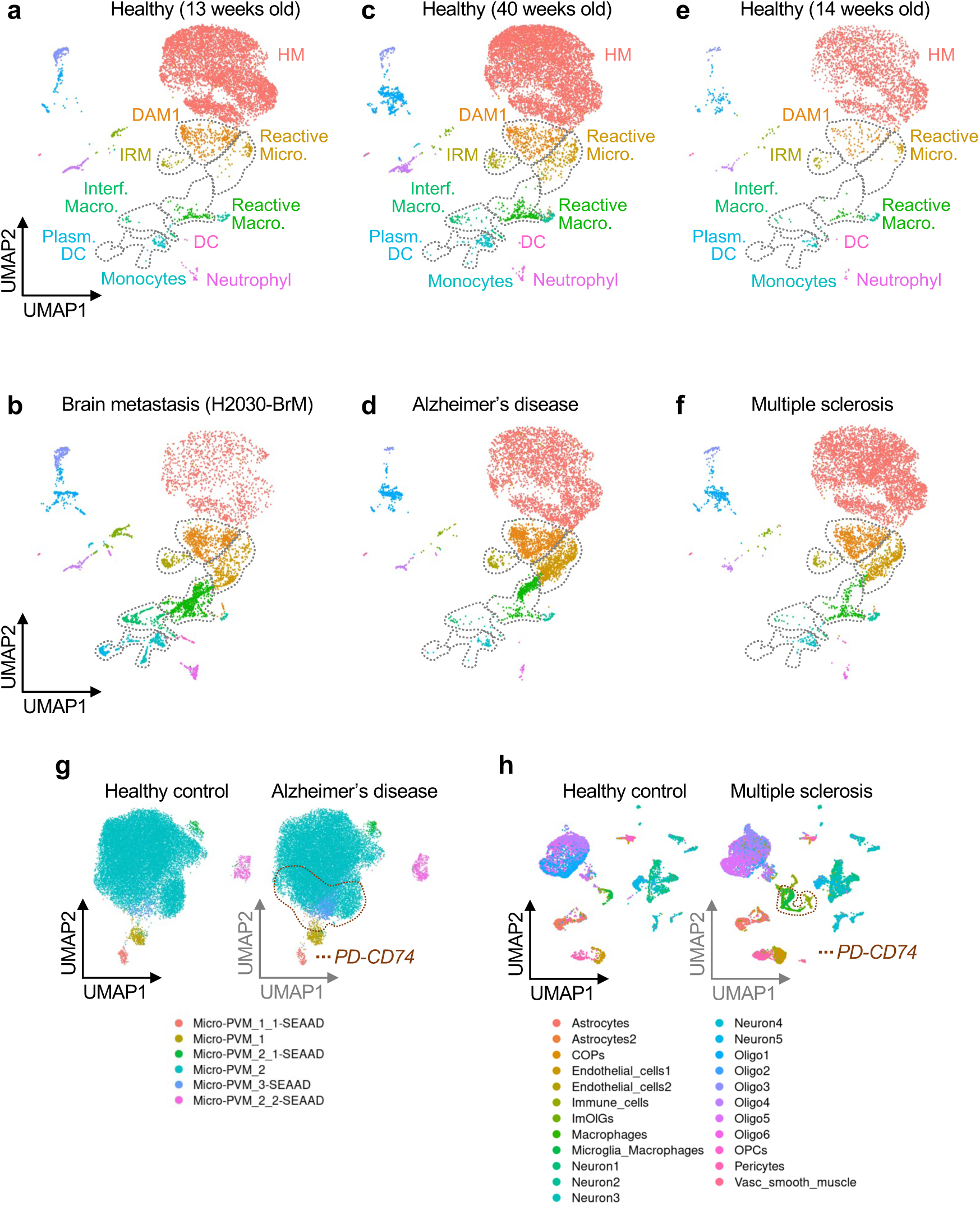
A CD74 pan-disease signature is present in Alzheimer’s disease and multiple sclerosis patients. **a. – f**. UMAPs of CD45^+^ populations found in ctrl healthy brain from immunocompromised mice (**a**), in H2030-BrM brain metastasis (**b**), in ctrl healthy aged brain from immunocompetent mice (**c**), in brain from Alzheimers disease mouse model (**d**), in ctrl healthy brain from immunocompetent mice (**e**), in brains from multiple sclerosis mouse model (**f**). HM: homeostatic microglia. DAM1: disease-associated microglia. IRM: interferon-responsive microglia. Reactive Micro.: reactive microglia. Reactive Macro: reactive macrophages. Interf. Macro.: interferon-responsive macrophages. DC: dendritic cells. Plasm. DC: plasmacytoid dendritic cells. **g.** UMAPs of cell type populations found in the dorsolateral prefrontal cortex of human brains from healthy controls and Alzheimer’s disease patients. **h.** UMAPs of cell type populations found in human brains from healthy controls and multiple sclerosis patients.

## Supplementary tables

**Table 1:** Brain metastasis samples analysed for MIF in either the tumor or the microenvironment. The % MIF was obtained by normalizing the area MIF^+^ for each compartment (either tumor or microenvironment) to the area of each compartment (either tumor or microenvironment) as determined by H&E.

**Table 2:** Differentially expressed genes in CD74^+^ vs CD74^−^ microglia/macrophages obtained by bulk RNA-seq. These genes include M1 and M2 markers, as well as tumor-associated microglia (TAM MG) and tumor-associated macrophages (TAM BMDM) markers that have been previously described in the literature.

**Table 3:** List of the top 150 up and 150 down differentially expressed genes in CD74^+^ vs CD74^−^ microglia/macrophages obtained by bulk RNA-seq.

**Table 4:** Differentially expressed genes in CD74^+^ CCR2^+^ cells vs CD74^+^ CCR2^−^ cells. The microglia/macrophage cluster was reanalyzed from GSE228368. Differentially expressed genes were obtained by comparing the *Cd74^+^ Ccr2^+^* populations to *Cd74^+^ Ccr2^−^* populations.

**Table 5:** Gene set enrichment analysis of CD74^+^ microglia/macrophages. Gene set enrichment analysis (GSEA) of top 25 upregulated and downregulated signatures in the bulk RNA sequencing of CD74^+^ microglia/macrophages compared to the CD74^−^ microglia/macrophages

**Table 6:** Differentially expressed genes for the effect of ibudilast in reactive macrophages from mice with brain metastasis (ibudilast vs. vehicle treatment).

**Table 7:** Patient-derived organotypic cultures (PDOC) treated with Ibudilast. Results from the Patient-derived organotypic brain cultures (PDOCs) of human brain metastasis treated with Ibudilast.

**Table 8:** Brain metastasis samples analyzed for CD74 in the microenvironment.

**Table 9:** MIF levels from ELISA applied to CSF from patient samples. Levels of MIF in the cerebrospinal fluid (CSF) of non-cancer patients and brain metastasis patients from different primary tumors.

**Table 10:** Upregulated pathways in the CD74^+^ myeloid clusters vs CD74^−^ myeloid clusters. Result of functional analysis highlighting the top upregulated pathways identified in the single cell RNA sequencing analysis of CD74^+^ myeloid clusters (reactive microglia, disease associated microglia, reactive macrophages) vs CD74^−^ myeloid clusters (homeostatic microglia, homeostatic macrophages) in brain metastasis, Alzheimer’s disease and multiple sclerosis.

**Table 11:** List of deregulated genes in the CD74^+^ microglia (disease associated microglia, reactive microglia)/macrophages (reactive macrophages) vs CD74^−^ microglia (homeostatic microglia)/macrophages (homeostatic macrophages) in brain metastasis, Alzheimer’s disease and multiple sclerosis.

## Notes

### Competing Interest Statement

The authors have declared no competing interest.

